# The aging modulator miR-29 is essential for adult cardiomyocyte function

**DOI:** 10.64898/2026.04.24.718944

**Authors:** David Roiz-Valle, Cintia Folgueira, Lucas Moledo-Nodar, Antonio G. Tartiere, Beatriz Cicuéndez, Rafael Romero-Becerra, Francisco Rodríguez, You-Wen He, José M.P. Freije, Guadalupe Sabio, Carlos López-Otín, Xurde M. Caravia, Alejandro P. Ugalde

## Abstract

Aging is the main risk factor for cardiovascular diseases, underscoring the need to identify the molecular regulators that sustain cardiac function during aging.

The microRNA miR-29 is a well-established aging-associated regulator as its expression increases with age, and its overexpression promotes premature aging. Here, we define the cardiomyocyte-autonomous role of miR-29 in the adult heart by generating an inducible, cardiomyocyte-specific miR-29-deficient mouse model (Heart-iKO). We show that Heart-iKO mice develop dilated cardiomyopathy (DCM) with reduced ejection fraction that ultimately leads to premature death.

Mechanistically, Heart-iKO cardiomyocytes exhibit alterations in mitochondrial structure and function. Transcriptomic profiling of bulk heart tissue and isolated cardiomyocytes revealed a consistent downregulation of genes involved in oxidative phosphorylation and the electron transport chain. We further observed a similar pattern of mitochondrial impairment in miR-29-deficient human cardiomyocytes derived from induced pluripotent stem cells (CM-iPSCs). Together, these findings highlight the context-dependent role of miR-29 in cardiac physiology and aging: although its upregulation promotes premature aging, its basal expression is required to maintain mitochondrial homeostasis and prevent heart failure in the adult myocardium.

## Introduction

Aging is the strongest risk factor for cardiovascular diseases (CVDs) (1). As the population ages, understanding the molecular pathways that preserve cardiac homeostasis becomes increasingly important. Cardiovascular aging contributes to heart failure, vascular dysfunction and mortality, and cardiovascular diseases remain the leading cause of death worldwide (2). In the last decade, the hallmarks of aging (3, 4), and more specifically of cardiovascular aging (5), have been established and reviewed several times. These advances have provided a framework for identifying molecular regulators that drive, or protect against, age-associated cardiac decline.

Among these regulators, microRNAs (miRNAs) have emerged as important post-transcriptional modulators of tissue homeostasis and stress responses. One representative group of geromodulatory microRNAs is the miR-29 family. This family comprises three members, miR-29a, miR-29b and miR-29c, encoded by two bicistronic genomic clusters (*miR-29a/b-1* and *miR-29b-2/c*) (4). miR-29 is highly conserved across species, its expression increases during both physiological and pathological aging (6, 7), and its overexpression promotes a premature aging phenotype (8). miR-29 dysregulation has also been implicated in several age-associated disorders, including cancer (9), metabolic diseases (10), and cardiovascular pathology (11).

In the cardiovascular system, miR-29 was initially linked to extracellular matrix remodeling after myocardial infarction (11). We previously showed that systemic loss of *miR-29a/b-1* in mice results in vascular remodeling, pulmonary congestion, and diastolic dysfunction, ultimately leading to heart failure with preserved ejection fraction (HFpEF) and premature death. This phenotype was accompanied by mitochondrial alterations in cardiomyocytes and was partially rescued by *Pgc1a* haploinsufficiency, pointing to altered mitochondrial homeostasis as a central mechanism (12).

However, the systemic and vascular abnormalities of these mice precluded distinguishing direct cardiomyocyte-specific effects from secondary consequences of whole-body miR-29 deficiency, suggesting a context-dependent role of miR-29 in tissue homeostasis. To address this limitation, we generated an inducible cardiomyocyte-specific miR-29 knockout model to define the cell-autonomous role of miR-29 in the adult myocardium. Here, we tested whether miR-29 is intrinsically required in cardiomyocytes to preserve mitochondrial homeostasis and cardiac function during adult life. We found that cardiomyocyte-specific loss of miR-29 is sufficient to disrupt mitochondrial homeostasis and drive dilated cardiomyopathy, leading to premature death.

## Materials and methods

### Animal care

All animal procedures listed in this study were previously approved by the Ethics Committee of the University of Oviedo and authorized by the Department of Rural Affairs and Agricultural Policy of the Principality of Asturias. Animals were housed in a pathogen-free animal facility under a 12-h day/12-h dark cycle, a temperature of 22 ± 2°C and a relative humidity of 50 ±10%, as well as *ad libitum* access to food and water.

### Mutant mice strain generation and genotyping

*miR-29^fl/fl^* mutant mice were generated and imported from the lab of Dr. You-Wen He (Duke University), and *Myh6-Cre* mutant mice (B6.FVB(129)-A1cfTg^(*Myh6-cre/Esr1\**)1Jmk^/J, Jackson #005657) were generated in the lab of Jeffery Molkentin (13), and obtained from the lab of Dr. Juan Miguel Redondo (Centro de Biología Molecular Severo Ochoa, CBMSO). To generate cardiomyocyte-specific miR-29 deficient mice (*Myh6-Cre miR-29^fl/fl^*, hereafter referred to as Heart-iKO), we crossed *Myh6-Cre* and *miR-29^fl/fl^*mouse lines. Heart-iKO mouse line was then backcrossed onto a C57BL/6N genetic background.

For genotyping, genomic DNA was isolated from mouse tail biopsies using an alkaline lysis buffer (NaOH 25 mM, EDTA 0.2 mM, pH 8) followed by a 99 °C incubation for 60 min and a subsequent addition of a neutralization buffer (Tris 40 mM, pH 7.4). 1 µl of lysate was used for PCR amplification using Platinum™ Taq DNA polymerase (Invitrogen™, #15966005) and specific primers for each mouse line (**Table 1**). Genotyping was performed by analyzing the size of the resulting PCR amplicons by electrophoresis in a 2% agarose gel.

**Table 1.**
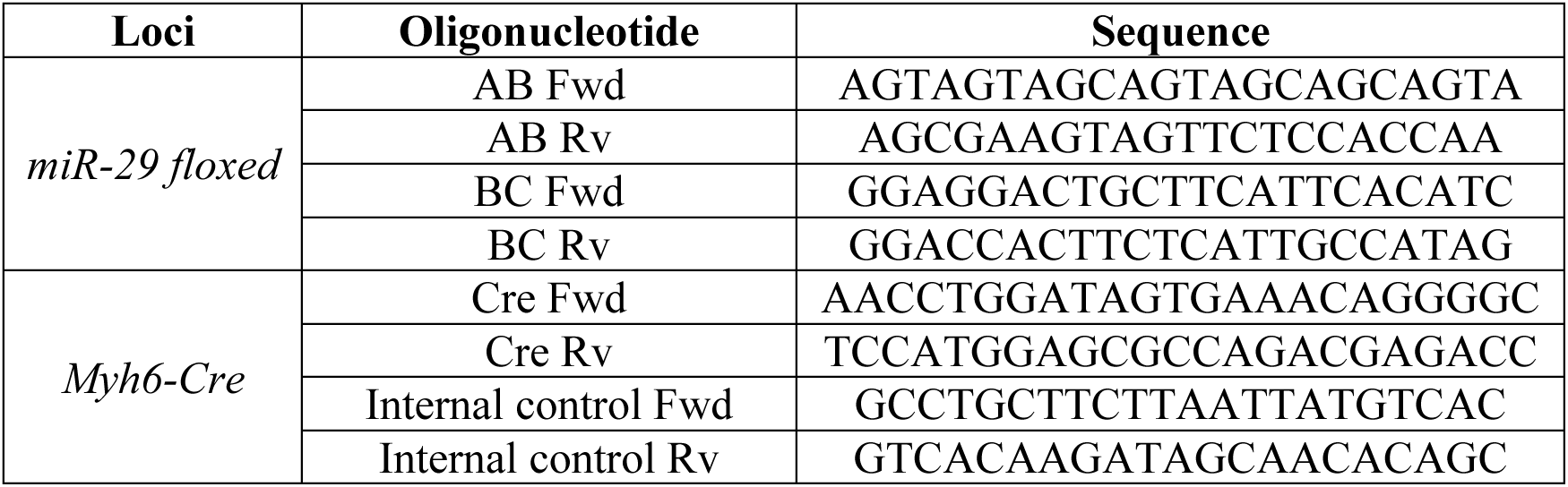
Oligonucleotides for PCR genotyping.

### Quantitative PCR (qPCR) of miR-29 expression in Heart-iKO mice

To verify the miR-29 expression in Heart-iKO mice, we harvested heart tissue samples, one month after 4-hydrotamoxifen treatment, from Heart-iKO mice and miR-29*^fl/fl^* controls. RNA was subsequently extracted using the TRIzol reagent protocol (Invitrogen™, #15596026). 10 ng of RNA from each sample were reverse-transcribed into miRNA cDNA using the TaqMan Advanced miRNA cDNA synthesis kit (Applied Biosystems, #A28007) and 1.5 µl of random primers (Invitrogen™, #48190011). qPCR was performed to relatively quantify the expression of each miR-29 family member in Heart-iKO samples compared to miR-29*^fl/fl^*controls. qPCR was performed in a QuantStudio 5 Real-Time PCR instrument using TaqMan Universal PCR Master Mix (Applied Biosystems, #4304437) and a predesigned TaqMan assay for each miRNA (Applied Biosystems # A25576, *478587_mir* for miR-29a, *478369_mir* for miR-29b and *479229_mir* for miR-29c). The endogenous housekeeping gene *18S* rRNA was used for normalization, and its levels were quantified using PowerUP™ SYBR™ Green Master Mix (ThermoFisher, #A25742) and the following oligonucleotides: 5’-CCATCCAATCGGTAGTAGCG-3’ (Fwd) and 5’-GTAACCCGTTGAACCCCATT-3’ (Rv).

To evaluate miR-29 expression directly in Heart-iKO cardiomyocytes, we purified adult mouse cardiomyocytes from seven-month-old mice. The isolation was performed in Centro Nacional de Investigaciones Cardiovasculares (CNIC) following the Langendorff protocol (14). Once isolated, cardiomyocytes were pelleted and frozen until further processing. RNA was extracted following TRIzol reagent protocol, and the RT and qPCR were performed as described above.

### Mitochondria isolation from Heart-iKO heart tissue and Oroboros respirometry

Heart-iKO and miR-29*^fl/fl^* mice were euthanized and their hearts were immediately harvested, cleaned in PBS and then kept on ice. Heart tissue samples were placed in a 100 µm filter on top of a 50 ml Falcon and washed with ice-cold PBS. The tissue was then placed in a tube along with 2 ml of Medium A (320 mM sucrose, 10 mM Tris-HCl, 1 mM EDTA, 0.1% fatty-acid-free BSA; pH 7.4) and homogenized using an automatic Potter-Elvehjem. Afterwards, the lysate was centrifuged at 1,000g at 4 °C for 5 min. The supernatant was then transferred to a new tube and centrifuged at 12,000g at 4 °C for 3 min. The supernatant was removed, and the resulting mitochondria pellet was resuspended in Medium A and centrifuged again at the same speed and time parameters. The process was repeated until the supernatant became clear and transparent. Then, the supernatant was removed, and the pellet was resuspended in buffer MAITE (25 mM sucrose, 75 mM sorbitol, 100 mM KCl, 0.05 mM EDTA, 5 mM MgCl_2_, 10 mM Tris-HCl, 10 mM K_2_HPO_4_, 0.1% fatty-acid-free BSA).

Activity of the electron transport chain (ETC) was assessed measuring oxygen consumption in a Clark type polarographic oxygen sensor (Oroboros instruments). Mitochondria were introduced inside a 2 ml isolated chamber, in agitation with a magnetic stirrer, at 37 °C, in MiR05 respiration medium (Oroboros instruments, # 60101-01). Once stabilized, they were sequentially incubated with different substrates and inhibitors: pyruvate (Sigma, #P2256) + malate (Sigma, #2300), ADP (Sigma, # A2754), rotenone (Sigma, #R8875), succinate (Sigma, #224731), antimycin A, *N*,*N*,*N*′,*N*′-Tetramethyl-*p*-phenylenediamine (TMPD, Sigma # T7394), and azide (Sigma, #S2002).

To analyze the measurements, respiration following antimycin A incubation was set as baseline. Then, complex IV activity was calculated as TMPD minus azide oxygen consumption rate, and all other measurements were normalized by this variable and by total µg of mitochondrial protein—measured using a Bradford assay. The calculations performed to obtain ADP-stimulated coupling and complexes I and II measurements were performed subtracting the OCR as indicated in the Oroboros instructions (**Table 2**).

**Table 2.** Mitochondrial respiration analysis from Oroboros instrument data.

| Variable | Calculation |
| --- | --- |
| Complex IV | TMPD - Azide |
| ADP-stimulated coupling | ADP - (Pyruvate + Malate) |
| Complex I | ADP - Rotenone |
| Complex II | Succinate - Antimycin A |

### iPSC culture

The HDF-iPS-SV10 human iPS cell line was obtained from the Spanish National Bank of Cell Lines. iPSCs were cultured with mTeSR™ Plus media (StemCell technologies, #100-0276) using Matrigel-coated cell culture plates. To coat the cell plates, 100 µl of Corning® Matrigel® Growth Factor Reduced Basement Membrane Matrix (Corning, #354230) were added to 6 ml of DMEM:F12 (1:1) + GlutaMAX™ media (Gibco, #10565018). The mixture was prepared on ice, and 1 ml was added to every well of a 6-well culture plate. Then, the plate was left for at least 1 h at 37 °C before seeding the cells.

iPSCs were routinely passaged using ReLeSR™ dissociation and selecting reagent (StemCell technologies, #100-0483), and frozen using mFreSR™ freezing media (StemCell technologies, #05855). During the first 24 h after passaging, mTeSR™ Plus media was supplemented with 10 µM Y-27632 (ROCK inhibitor, StemCell technologies, #72304).

### CRISPR editing of human iPSCs

To generate a *miR-29a/b-1*-deficient iPSC line, HDF-iPS-SV10 human iPSCs were edited using the CRISPRevolution sgRNA EZ Kit (Synthego) in the Pluripotent Cell Technology technical unit of CNIC using custom designed sgRNA sequences: mir-29a, 5’-GGAAATGTATTGGTGACCGT-3’; miR-29b-1, 5’-CTTAAGAATAAGGGAGTCCC-3’.

To prepare the ribonucleoprotein (RNP) complex mix, sgRNAs were incubated at 95 °C for 5 min. Then, 0.75 µl of 10 µg/µl Cas9-HIFi v3 (IDT technologies, # 1081060) were mixed with the sgRNAs and the mixture was incubated at room temperature for 20 min. The RNP mix and 1 µg of pEGFP plasmid (Addgene, #165830) were then added to 1 million iPSCs and transfected using electroporation. To this end, a NEON electroporation system was used, with 1 pulse of 30 ms at 1,200 V. Following electroporation, cells were plated in a 24-well plate in iPS-Brew medium (Miltenyi Biotec, #170-076-317) supplemented with 10 µM ROCK inhibitor. After 48 h, cells were dissociated using accutase (ThermoFisher, #A1110501) and sorted based on GFP expression. GFP-positive and GFP-negative populations were plated separately and maintained in iPS-Brew medium with 10 µM ROCK inhibitor. The medium was replaced 48 h later with inhibitor-free medium, and the cells were cultured for three additional days. Subsequently, cells were dissociated again, plated at low density on 100 mm dishes, and cultured until individual colonies were large enough for picking. During this expansion period, the medium was changed daily. Individual colonies were then picked and expanded. Before cryopreservation, a replicate of each clone was lysed for genomic DNA extraction, which was then screened by PCR to confirm the presence of the mutation. Five confirmed clones were selected for further experimentation as *miR-29a/b-1*-deficient human iPSCs.

### Generation of cardiomyocytes derived from human iPSCs (CMs-iPSC)

Highly confluent iPSCs were differentiated into cardiomyocytes (CM-iPSCs) using a protocol involving time-controlled supplementation of differentiation factors (**Table 3**) (15). At day 0 of differentiation, Basal[-] media plus 10 µM Chir (Sigma, #SML1046) was added to the cells. After 24 h, the media was removed and exchanged for Basal[-]. At day 3, media was replaced for Basal[-] supplemented with 5 µM WNT-C59 (Sigma, #5004960001). At day 5, media was again replaced for Basal[-] and left for two days. At day 7, media was exchanged for Basal[+]. After 3 days, metabolic selection was started by replacing the media with Selective media, which only contains lactate as fuel source and selects differentiated cardiomyocytes. Selective media was replaced every two days until non-cardiomyocyte cells were eliminated (CM-iPSCs can be distinguished by their autonomous beating). After that, media was replaced for Basal[+] and differentiated CM-iPSCs were then left for a week maturing before replating for downstream experiments. To replate CM-iPSCs, cells were detached from the plate by incubating them with Gibco™ TrypLE™ Express (Gibco, #10043382) for 15 min at 37 °C. After seeding at the desired concentration, cells were left untouched for 2-3 days with Basal[+] media supplemented with 10% KnockOut™ Serum Replacement (Gibco, #10828028), 10 µM ROCK inhibitor and 2 µM Chir.

**Table 3.** Media needed for CM-iPSC differentiation.

| Media | Component | Quantity |
| --- | --- | --- |
| **Basal minus Insulin ([-])** | RPMI1640 + GlutaMAX™ | 500 ml |
|  | Ascorbic acid | 500 µl |
|  | B27 supplement minus insulin | 10 ml |
| **Basal ([+])** | RPMI1640 + GlutaMAX™ | 500 ml |
|  | Ascorbic acid | 500 µl |
|  | B27 supplement (with insulin) | 10 ml |
| **Selective** | RPMI1640 minus glucose | 500 ml |
|  | CDM3 supplement | 5 ml |
|  | Sodium lactate | 420 µl |
| **CDM3 supplement** | L-ascorbic acid 2-phosphate | 4.224 g |
|  | Recombinant human serum albumin | 10 g |
|  | H<sub>2</sub>O | 200 ml |
| Ascorbic acid (Sigma, #A92902), B27 minus insulin (Gibco, #A1895601), B27 with insulin (Gibco, #17504044), sodium lactate (Sigma, #L4263), L-ascorbic acid 2-phosphate (Sigma, #A8960), recombinant HSA (Sigma, #A9731). |  |  |

### Seahorse Mito Stress Test

To assess mitochondrial function in CM-iPSCs, we used a Seahorse XF HS mini instrument, adapting an already-described protocol (16). Briefly, XFp miniplates were coated with Matrigel as previously described and 10,000 cells per well were seeded. Basal[+] culture medium was changed for Seahorse XF RPMI Basal medium (Agilent, #103576-100) supplemented with 2 mM glutamine, 10 mM glucose and 1 mM sodium pyruvate 1 h before the start of the assay, leaving the cells in a 37 °C non-CO_2_ incubator until the beginning of the assay. Following Seahorse Mito Stress assay standard protocol, three different combinations of compounds were sequentially injected during the assay: 2.5 µM oligomycin, 2 µM FCCP and 2.5 µM rotenone/antimycin A. After the assay, cells were fixed with 4% PFA, permeabilized with 1% Triton X-100 and then the total number of cells for each well was obtained by DAPI staining and subsequent nuclei count of fluorescence microscope images from each well. That number was then used for normalization of Seahorse data.

### Cre recombinase induction with 4-OH tamoxifen

Heart-iKO and control *miR-29^fl/fl^* mice were treated at 8 weeks of age with 4-hydroxytamoxifen (4-OH TX) (Selleckchem, #S7827). 1 mg of 4-OH TX was administered intraperitoneally each day during 5 consecutive days, previously solubilized in 90% corn oil (Sigma, #C8267) and 10% ethanol. Control vehicle was prepared with the same proportion of corn oil and ethanol but without tamoxifen.

### Blood pressure and heart rate analysis of Heart-iKO mice

Blood pressure and heart rate in mice were measured using the non-invasive tail-cuff method (17). The procedure was performed in conscious mice placed in a BP-2000 Blood Pressure Analysis System (Visitech Systems). 10 preliminary measurements and 10 experimental measurements were recorded, and the average of the 10 experimental measurements was used for analysis. The animals were trained for four consecutive days prior to recording the measurements for analysis. All the procedures were performed at the same time of the day.

### Electrocardiogram analysis

ECG recordings were acquired at 2 kHz using a MP36R data acquisition workstation (Biopac Systems) and exported with *AcqKnowledge* software (Biopac Systems) for automatic analysis with custom R and Matlab scripts, as described in (18).

### Echocardiography

Heart-iKO and *miR-29^fl/fl^* mice were anesthetized by inhalation of isoflurane, and echocardiography was performed with a 30 MHz transthoracic echocardiography probe. Images were obtained with a Vevo 2100 micro-ultrasound imaging system (VisualSonics, Toronto, Canada). Short-axis, long-axis, B-mode, and two-dimensional M-mode views were obtained. Scans were conducted by two experienced researchers blinded to the mouse genotypes. Measurements of left parasternal long and short axes and M-mode images (left parasternal short-axis) were obtained at a heart rate of 500-550 beats per min. Left ventricle (LV) end-diastolic diameter (LVEDD), LV end-systolic diameter (LVESD), and wall thickness were measured from M-mode tracings, and the average of three consecutive cardiac cycles was reported. Systolic cardiac function was estimated from the ejection fraction (EF) and fractional shortening (FS) obtained from M-mode views. The LV fractional shortening percentage was calculated as *([LVEDD - LVESD] / LVEDD) × 100*. For ejection fraction measurements, a long- or short-axis view of the heart was selected to obtain an M-mode registration in a line perpendicular to the left ventricular septum and posterior wall at the level of the mitral *chordae tendineae*.

### Histological procedures

Heart-iKO and *miR-29^fl/fl^* mice were euthanized and their hearts were then washed in PBS, weighted, and immediately immersed in Krebs Henseleit cardiac relaxant solution (118 mM NaCl, 4.7 mM KCl, 1.2 mM KH_2_PO_4_, 1.2 mM MgSO_4_, 25 mM NaHCO_3_, 11 mM glucose). After 30 min, hearts were transferred to a paraformaldehyde (PFA) 4% in PBS solution and stored at 4 °C during 48 h. Later, PFA was changed for 50% ethanol, where they were kept until inclusion. Paraffin embedding of PFA-fixated samples, as well as tissue sectioning and staining with hematoxylin and eosin (H&E) and Massońs trichrome, was performed by the molecular histopathology facility of the Instituto de Oncología del Principado de Asturias (IUOPA).

For WGA staining, paraffin-embedded hearts were rehydrated in the molecular histopathology facility and then washed 3 times with HBSS 1X with Ca^2+^ and Mg^2+^ (10X stock from Gibco™, #14065056). Then, tissue sections were subsequently blocked with 5% BSA in TBS-T, and incubated with WGA-CF®488A (Biotium, #29022) for 2 h. Afterwards, green channel fluorescence of the samples was imaged in a Zeiss Axio Observer microscope under a Plan-Neofluar 20x/0.50 Ph 2 objective.

Heart and lung tissue images were analyzed using *QuPath*. For H&E images, a pixel thresholder was used to detect tissue area, and custom scripting was used to extract the filled area (adding holes), and from it calculate hole area. For Masson trichrome images, first stain vectors were set selecting collagen-stained areas, and then pixel thresholders were used to detect subsequently full tissue area and collagen-occupied tissue area. *Fiji/ImageJ* was also used for image processing to generate figure panels.

### Ultrastructural analysis with transmission electron microscopy

Heart-iKO and *miR-29^fl/fl^* mice were euthanized and their hearts were immediately cleaned in PBS and introduced in a fixative solution made of 4% PFA and 2.5% glutaraldehyde in PBS. The hearts were cut off in cubical sections of approximately 1 mm^3^, avoiding tearing the tissue during the cutting procedure. Each cubical section was then placed in an eppendorf tube with 1 ml of fixative solution and left 2-3 h at room temperature. After that time, fixative solution was changed for fresh one, and samples were left overnight at 4 °C. The next day, the fixative solution was replaced with PBS and the samples were stored at 4 °C until processing. Inclusion, slicing, staining, and imaging of the samples was performed in the electron microscopy facility of the Spanish National Biotechnology Center (CNB-CSIC). From each individual analyzed, three 2,000x and eight 4,000x images were taken.

To analyze mitochondrial ultrastructure parameters, the images obtained from TEM were processed using *MitoNet*, a deep learning model to segment mitochondria, in the form of a *napari* plugin called *empanada* (19). Once segmented, the *scikit-image* Python module was used to extract the properties from every detected mitochondrion (area, major and minor axis lengths, perimeter and total image area). From those values, we calculated the circularity *(4*π*(area/perimeter^2^))*, aspect ratio *(major axis length / minor axis length)* and roundness *(4 * (area/(π* major axis length^2^)))*. Statistical analysis of the obtained data was performed in R. Plotting was performed using the package *tidyplots* (20).

### Transcriptomic analysis of miR-29-deficient mice

Total RNA was extracted from multiple tissues as previously described. All groups were euthanized and samples were extracted in the morning. The extracted RNA was then used for transcriptome sequencing by RNA-Seq. Library preparation and sequencing for Heart-iKO RNA was conducted by Macrogen using TruSeq stranded mRNA library and NovaSeq 6000 sequencing system generating 40 million reads per sample, paired-end and 150 bp length. FASTQ quality was checked using the *FASTQC* software (Babraham Bioinformatics) and adaptors were trimmed using *cutadapt* version 4.4. Gene expression was later quantified from the FASTQ samples using Salmon version 1.10.3. As a reference transcriptome, we used GENCODE GRCm39 M32. Differential expression analysis was performed using *DESeq2* (21).

To perform gene set enrichment analysis (GSEA), a ranking variable was created as follows: *rank = -log_10_(p-value) * sign(log_2_ Fold Change)*. The ranked vector of genes was compared to the Hallmarks and Gene Ontology collections from Mouse Molecular Signatures database (22). GSEA was performed using the *fgsea* package (23).

### Statistical analysis

To compare differences between two groups, parametric two-tailed Student’s *t* test was performed in data with normal distribution and homoscedasticity (previously checked with a Shapiro-Wilk and an F-test, respectively). For non-parametric data, a Mann-Whitney test was performed. All comparisons between two groups were done using the R package *tidyplots*. In relative expression analysis, two-way ANOVA (*average Ct ∼ Gene*Genotype*) and a subsequent Tukey post-hoc analysis was performed using R *stats* core library. For weight progression, data was analyzed by fitting a linear mixed-effects model (*Weight ∼ Group * Sex * Age + (1 | Mouse ID))* with the R *lmer* package and p-values were calculated using Satterthwaite’s method with the R *lmerTest* package (24). To compare the proportion of aberrantly large and aberrantly shaped mitochondria, observations were organized in contingency tables, and a chi-squared test was performed. For survival comparisons, the Log-rank test was performed using the *survival* R package (25). Statistical significance was defined as p < 0.05.

### Data and code availability

Raw data from transcriptomic analyses can be found on the European Nucleotide Archive with accession code PRJEB105187. All custom code created for these analyses is available at https://github.com/GenCanAgeLab/miR29-HeartiKO.

## Results

### Adult cardiomyocyte-specific deletion of miR-29 leads to premature death in mice

To characterize the role of miR-29 in adult cardiomyocyte function, we generated an inducible, cardiomyocyte-specific miR-29 deficient model (*Myh6/Cre^Tg+^ miR-29^fl/fl^*, Heart-iKO) by crossing miR-29 floxed (*miR-29^fl/fl^*) mice, which have both miR-29 clusters flanked by loxP regions, with the *Myh6-MerCreMer* mouse strain, which expresses an inducible Cre recombinase under the control of the cardiomyocyte-specific alpha myosin heavy chain (*Myh6*) gene promoter. To induce miR-29 deletion, both Heart-iKO mice and control *miR-29^fl/fl^*littermates were treated intraperitoneally for 5 consecutive days with 1 mg of 4-OH tamoxifen at 8 weeks of age.

Heart-iKO mice appeared phenotypically indistinguishable from their *miR-29^fl/fl^* littermates, with no overt alterations or changes in body weight (**Fig. 1a,b**). RT-qPCR analysis of bulk heart RNA revealed that miR-29 levels tended to decrease only in tamoxifen-treated Heart-iKO mice, thereby confirming that Cre recombinase remained inactive in the absence of tamoxifen. Nevertheless, the reduction was moderate and non-significant **(Suppl. Fig. 1a)**. However, analysis of miR-29 expression in RNA from isolated Heart-iKO cardiomyocytes revealed an almost complete suppression of miR-29 expression in these cells (**Fig. 1c**). This disparity can be explained by the fact that cardiomyocyte gene expression represents only a fraction of the total bulk heart transcriptome (26).

**Figure 1.**
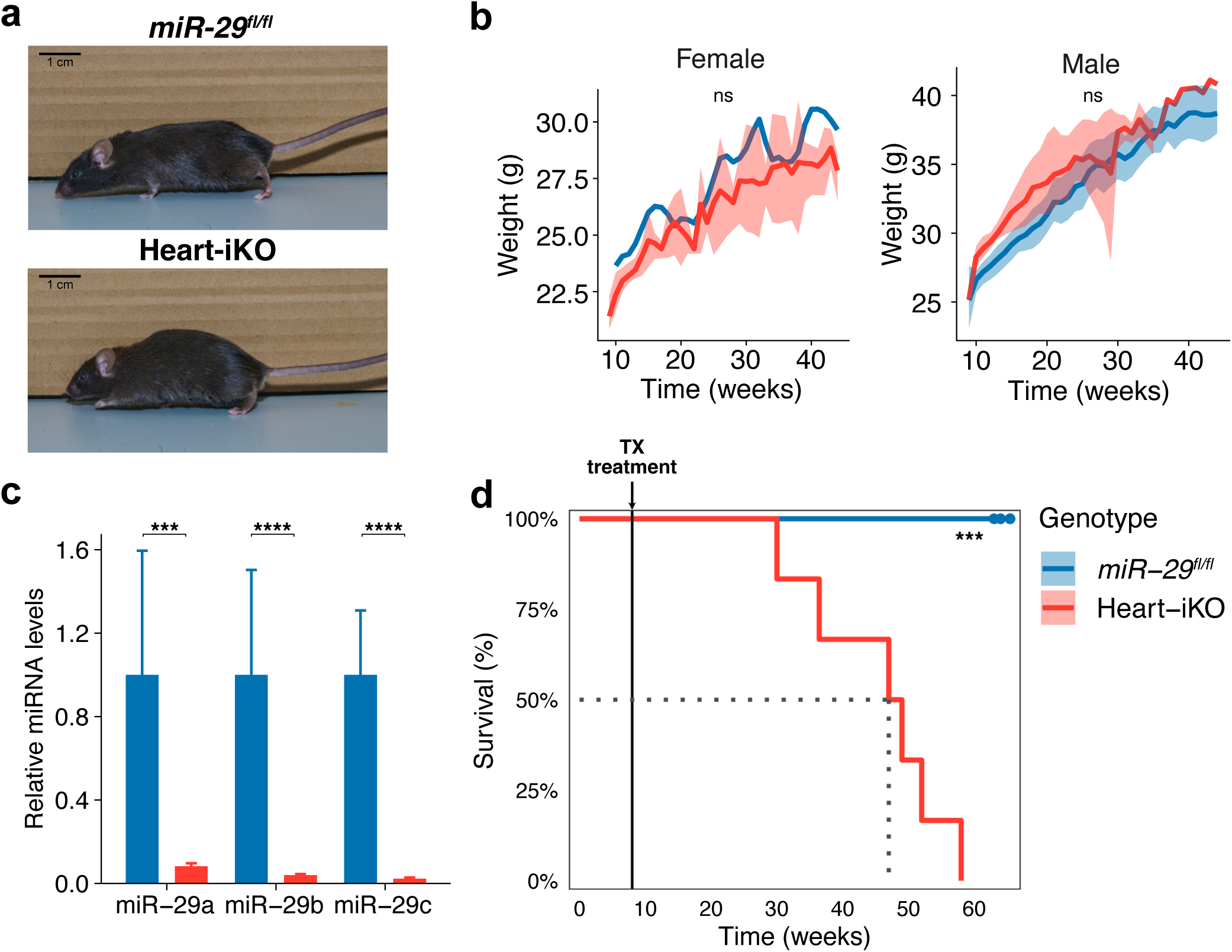
Characterization of Heart-iKO mice. a) Representative picture of *miR-29^fl/fl^* and Heart-iKO at 5 months of age. b) Weight progression (n=6 male and 1 female *miR-29^fl/fl^*, and 3 male and 3 female Heart-iKO mice). c) Relative expression of miR-29 family members in cardiomyocytes isolated from *miR-29^fl/fl^* (n=2) and Heart-iKO (n=3) mice at 5 months of age. d) Survival curve of *miR-29^fl/fl^* and Heart-iKO mice (n=6 male and 1 female *miR-29^fl/fl^* and 3 male and 3 female Heart-iKO mice). Bar and weight line plots represent mean ± SEM. Statistical significance in the weight progression data was determined by adjusting to a linear mixed-effects model (*Weight ∼ Group * Sex * Age + (1 | Mouse ID))* and then calculating the p-value for *Group*, *Group*Sex* and *Group*Sex*Time* using Satterthwaite’s method (all were *ns*). In the relative expression data, statistical significance was calculated using a two-way ANOVA and a post-hoc Tukey analysis. In the survival analysis data, statistical significance was obtained using the log-rank test. *: p < 0.05, **: p < 0.01, ***: p < 0.001, ****: p < 0.0001, ns: p > 0.05.

Despite the absence of overt phenotypic differences from *miR-29^fl/fl^*controls, Heart-iKO mice exhibited a drastically reduced lifespan, with a median survival of approximately 50 weeks (**Fig. 1d**). To rule out the presence of nonspecific Cre expression and to ensure that *miR-29^fl/fl^* mice did not exhibit alterations due to their genetic construction, we also monitored Heart-iKO and *miR-29^fl/fl^*mice treated with control vehicle (corn oil) instead of tamoxifen. Neither group, including tamoxifen-treated *miR-29^fl/fl^* mice, showed any apparent reduction in survival **(Suppl. Fig. 1b)**. Furthermore, there were no significant differences between male and female Heart-iKO survival curves, although male Heart-iKO mice showed a slightly shorter median survival than females **(Suppl. Fig. 1c)**.

### Heart-iKO mice present reduced systolic function and DCM

To better define the cardiovascular alterations that potentially lead to the premature death of Heart-iKO mice, we performed a comprehensive cardiovascular characterization. First, we measured blood pressure non-invasively in live animals. Previous studies reported a systemic hypertensive phenotype in whole body *miR-29a/b-1*-deficient mice (12). However, Heart-iKO mice showed no significant alterations either in systolic or in diastolic pressures **(Suppl. Fig. 2a,b)**. Focusing on the characterization of the cardiac function, we measured heart rate in Heart-iKO mice and found no significant differences compared to control mice **(Suppl. Fig. 2c)**. Moreover, an electrocardiogram analysis revealed no electrical abnormalities, as main electrical cardiac parameters—such as RR, which reflects heart rate; corrected QT interval (QTc), an indicator of arrhythmias; or QRS, which represents ventricular depolarization—were unchanged compared with *miR-29^fl/fl^*littermates **(Suppl. Fig. 2d-f)**.

To further study Heart-iKO heart physiology, we assessed cardiac structure and function through an echocardiography study. We found no alterations in heart wall thickness parameters, such as left ventricle posterior wall (LVPW) and intraventricular septum (IVS) thickness (**Fig. 2a, Suppl. Fig. 2g)**. In addition, diastolic function parameters, such as isovolumetric relaxation time (IVRT) or E/A ratio (**Fig. 2b,c**), were like those of *miR-29^fl/fl^* littermates. However, systolic function parameters, such as EF and FS, were significantly decreased compared with *miR-29^fl/fl^* control mice (**Fig. 2d,e**). Moreover, heart lumen—measured as left ventricle diameter in diastole—was significantly increased (**Fig. 2f**). These results indicated reduced heart contraction resulting in a systolic dysfunction and heart failure phenotype in Heart-iKO mice (**Fig. 2g**).

**Figure 2.**
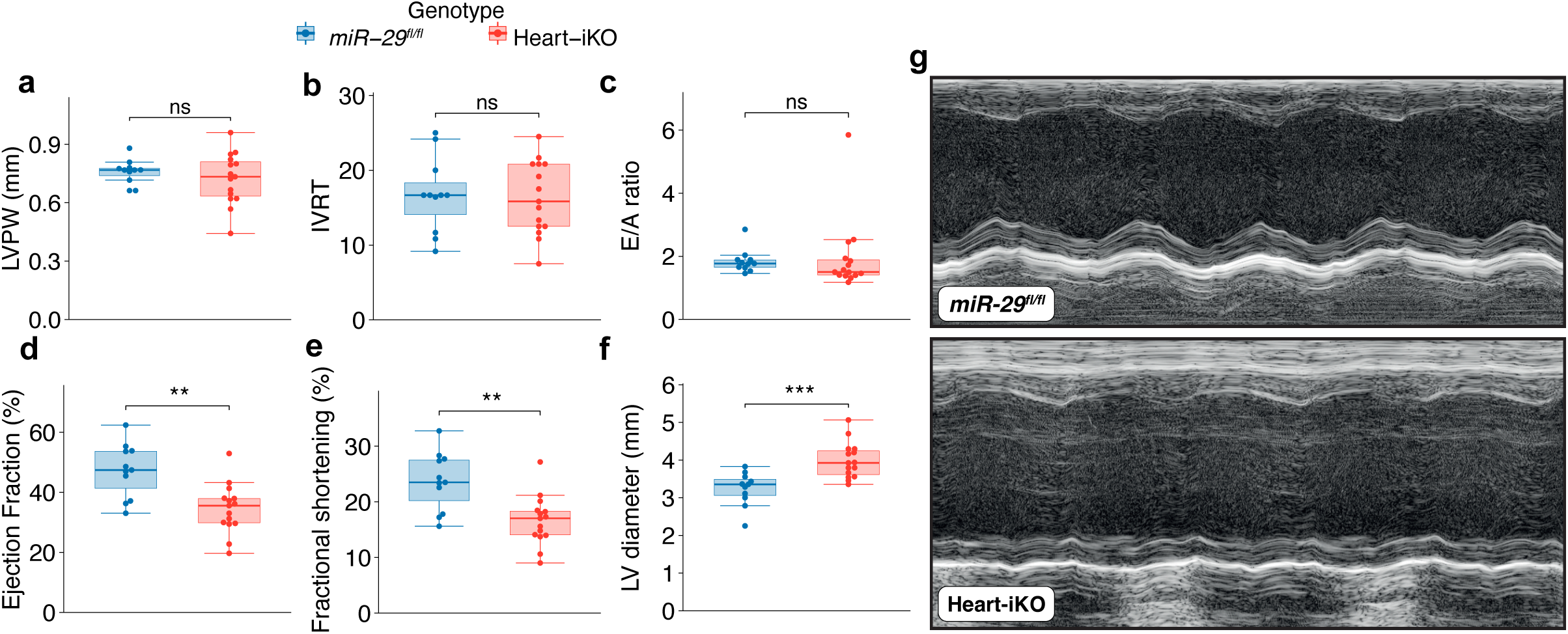
Echocardiography analysis of Heart-iKO mice. a) Left ventricle posterior wall (LVPW) thickness, b) isovolumetric relaxation time (IVRT), c) E/A ratio, d) ejection fraction (EF), e) fractional shortening (FS), f) left ventricle (LV) diameter measured in diastole, and g) Representative M-mode echocardiography. A total of 15 Heart-iKO and 11 *miR-29^fl/fl^*5-month-old male mice were studied. Statistical significance was determined using a non-parametric Wilcoxon signed-rank test. *: p < 0.05, **: p < 0.01, ***: p < 0.001, ****: p < 0.0001, ns: p > 0.05.

To confirm the alterations found in the echocardiography, we performed a morphological and histological study of Heart-iKO mouse heart. Heart-iKO mouse hearts were macroscopically bigger and weighed significantly more than *miR-29^fl/fl^* control mice (**Fig. 3a**). Histological results further supported the increased size of Heart-iKO heart, together with an increased lumen (**Fig. 3b,c**). Overall, echocardiographic, morphological and histological data indicate impaired systolic function of Heart-iKO mice, consistent with DCM phenotype. In line with this, most dilated Heart-iKO hearts also displayed characteristic symptoms of DCM such as cardiomyocyte disorganization **(Suppl. Fig. 3a)** and pulmonary edema **(Suppl. Fig. 3b)**. Furthermore, Heart-iKO hearts presented significantly increased accumulation of interstitial collagen fibers, which is consistent with the heart failure phenotype (**Fig. 3d**).

**Figure 3.**
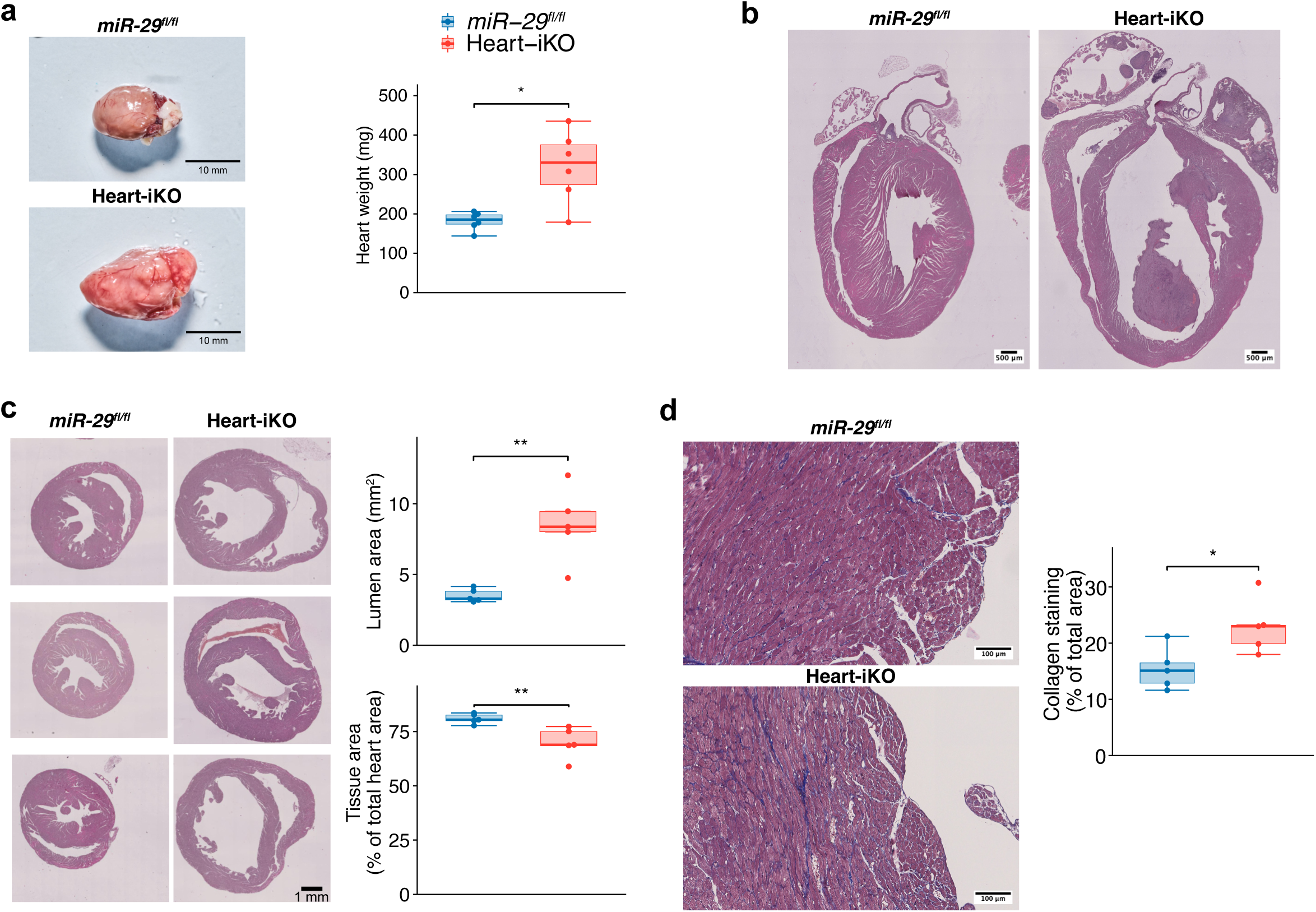
Cardiac morphology and histology of Heart-iKO mice. Morphological and histological analysis of 6-month-old Heart-iKO and *miR-29^fl/fl^* mice. a) Representative macroscopic images and weight of heart (n=6 per group). b) Representative H/E-stained 4-chamber view histology. c) Representative transversal sections, and lumen area and relative tissue area measurements (n=5 per group). d) Representative Masson trichrome staining and quantification of collagen-occupied area (n=5 per group). Statistical significance was determined using a non-parametric Wilcoxon signed-rank test. *: p < 0.05, **: p < 0.01, ***: p < 0.001, ****: p < 0.0001, ns: p > 0.05.

### Cardiomyocyte mitochondria in Heart-iKO mice exhibit structural and functional abnormalities

After identifying DCM in Heart-iKO mice, we investigated the potential causes of this dysfunction at the cellular level. To this end, we performed an ultrastructural study using TEM to identify alterations in cardiomyocytes (**Fig. 4a**). In previous studies using whole-body *miR-29a/b-1^-/-^* mice, we found that the loss of miR-29 induced changes in both abundance and structure of cardiomyocyte mitochondria. Therefore, we focused on mitochondrial morphology in TEM images. Although we did not find changes in mean mitochondrial size **(Suppl. Fig. 4a)**, we observed that the circularity of mitochondria in Heart-iKO cardiomyocytes was significantly reduced compared to *miR-29^fl/fl^* (**Fig. 4b**). Additionally, there was a decrease in the number of mitochondria per area, as well as in the percentage of area occupied by mitochondria (**Fig. 4c,d**). Furthermore, the percentage of mitochondria with unusually large sizes or aberrant shapes was increased in Heart-iKO cardiomyocytes **(Suppl. Fig. 4b, Fig. 4e**). Together, these results reveal marked alterations in mitochondrial organization in Heart-iKO cardiomyocytes.

**Figure 4.**
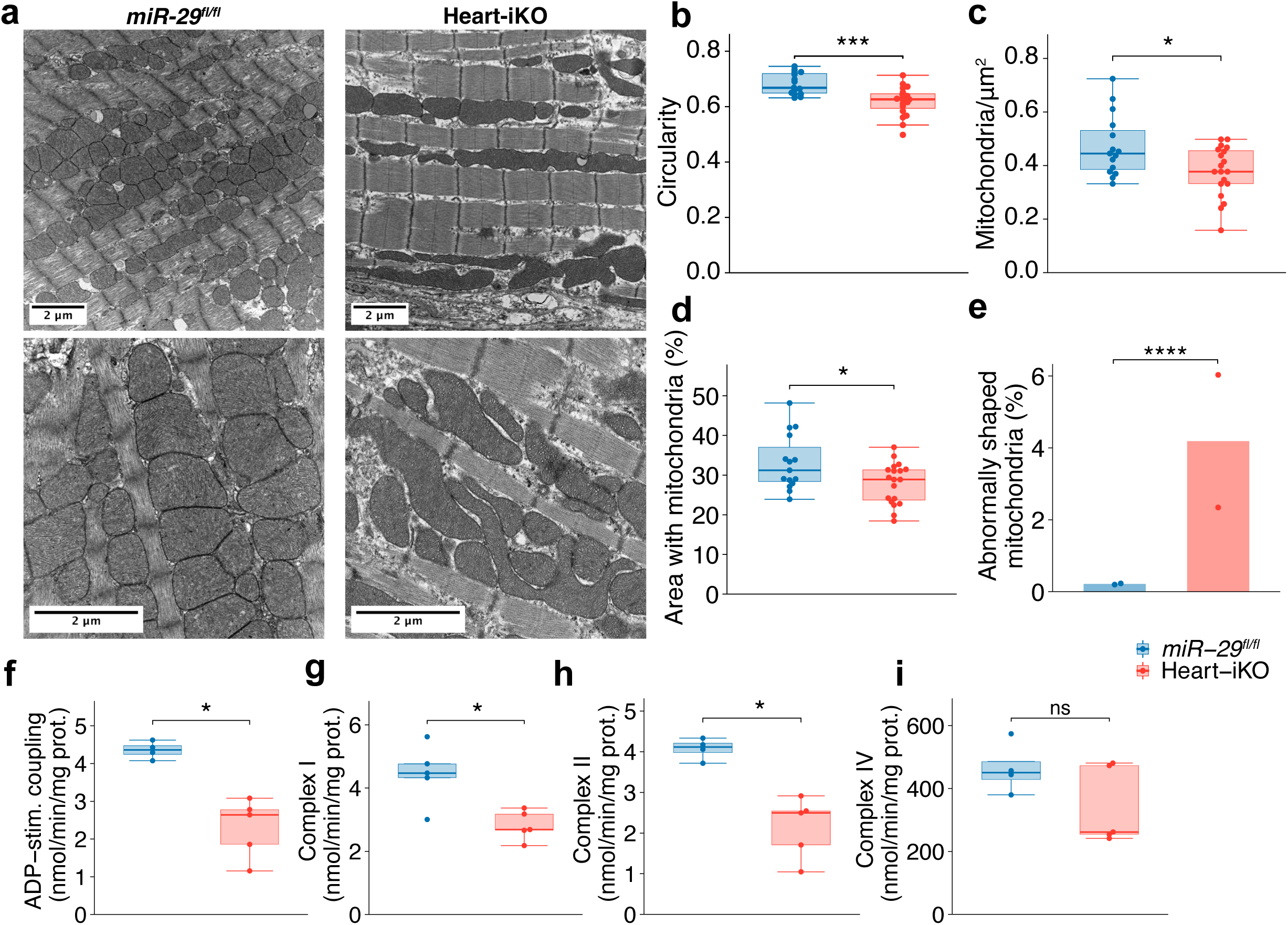
Mitochondrial structure and function in Heart-iKO. a) Representative transmission electron microscopy (TEM) images of 5-month-old *miR-29^fl/fl^* and Heart-iKO mice. b) Circularity of mitochondria. c) Mitochondrial density (mitochondria per µm^2^). d) Percentage of cardiomyocyte area occupied by mitochondria. e) Percentage of mitochondria with abnormal shape (circularity < 0.3). For TEM analysis, images were obtained from 5 tissue slices from 2 male mice per genotype. Each data point represents one image (n=15 *miR-29^fl/fl^*, 19 Heart-iKO). Images were obtained from 5 different tissue slices from 2 different mice per genotype. f-i) Mitochondrial activity measured using the Oroboros instrument: f) ADP-stimulated coupling activity, g) activity of complex I, h) activity of complex II, and i) activity of complex IV in mitochondrial extracts from heart tissue of 5-month-old male Heart-iKO and *miR-29^fl/fl^* mice (n=6 per group). Bar plots represent mean values. Statistical significance was determined using a non-parametric Wilcoxon signed-rank test, except for the percentage of abnormally shaped mitochondria, where data was structured as a contingency table and analyzed using a chi-squared test. *: p < 0.05, **: p < 0.01, ***: p < 0.001, ****: p < 0.0001, ns: p > 0.05.

To further study mitochondrial function, we studied these organelles using the Oroboros instrument, a high-resolution respirometry platform. We isolated mitochondria from Heart-iKO and *miR-29^fl/fl^* hearts and measured oxygen consumption changes in the extracts after supplementation with different activators and inhibitors of the mitochondrial ETC. We evaluated the activity of the ETC, as well as the individual activities of complexes I, II, and IV. We found that global OxPhos capacity was substantially reduced in Heart-iKO heart mitochondria, as well as complex I and complex II specific activities (**Fig. 4f-h**). However, there were no alterations in complex IV activity, which is known to possess excess capacity under ADP and oxygen saturation levels (27) (**Fig. 4i**). These results indicate impaired OxPhos-dependent ATP production in Heart-iKO mice, affecting both NADH-driven (Complex I) and FADH_2_-driven (Complex II) pathways.

### Heart-iKO transcriptomics reveal changes in genes implicated in OxPhos

Once we established that Heart-iKO mice exhibited DCM and dysfunctional mitochondria, we next examined the molecular alterations underlying this phenotype. To this end, we performed an RNA-Seq study of bulk cardiac RNA from 3-month-old Heart-iKO and *miR-29^fl/fl^* mice. A differential expression analysis identified 135 significantly upregulated and 60 significantly downregulated genes in Heart-iKO transcriptome (**Fig. 5a**). We cross-checked our gene list with data from TargetScan, miRDB and microT databases, which contain computational predictions regarding miRNA:mRNA interactions. Thus, among the top upregulated genes in our experiment, we found multiple predicted miR-29 targets, such as *Col4a5*, *Serpinh1*, *Col4a2*, *Narf*, *Dnmt3a*, and *Bdh1* (**Table 4**).

**Figure 5.**
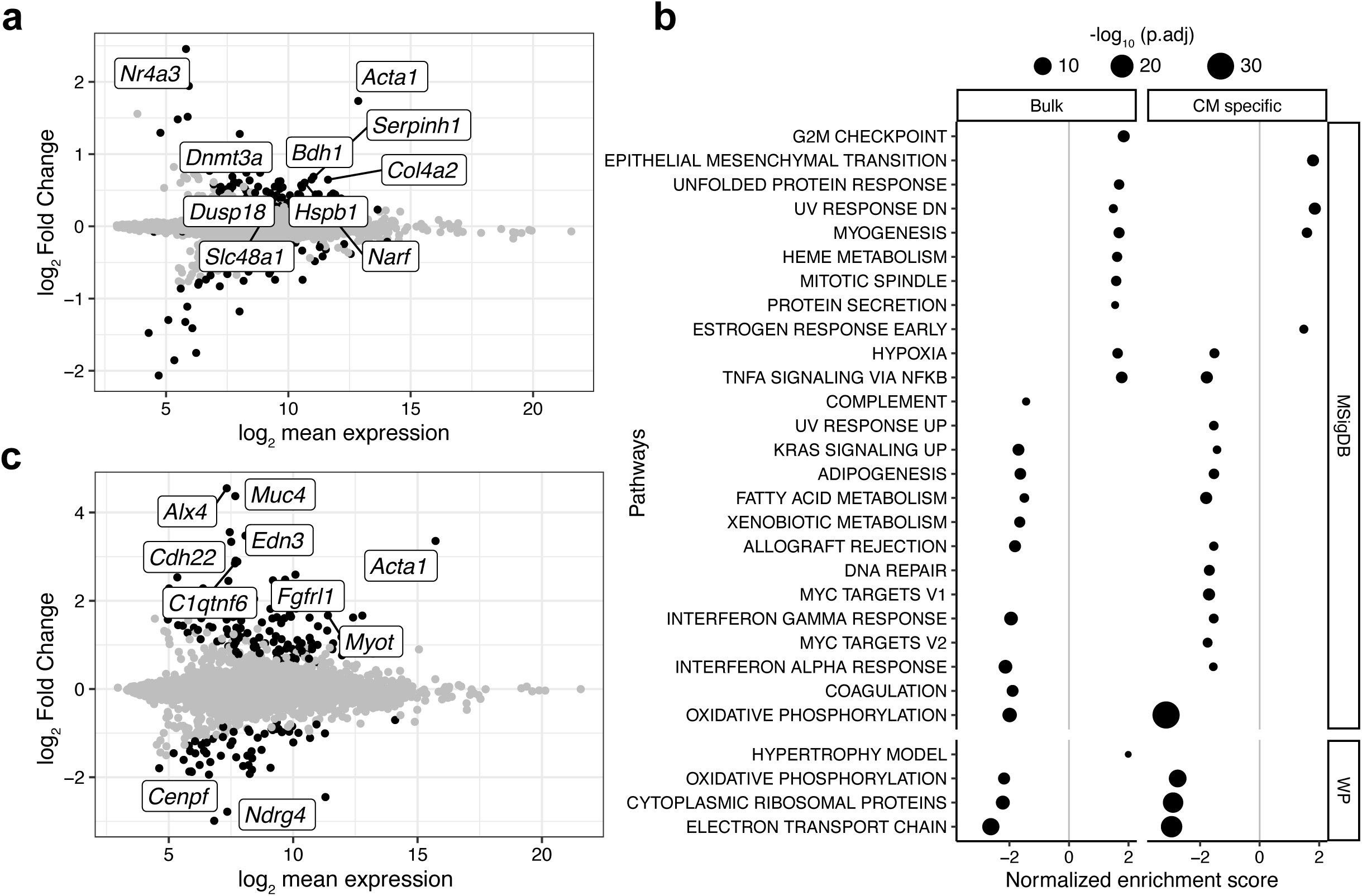
Heart-iKO transcriptomic analyses. a) MA plot displaying significant differentially expressed genes in bulk heart transcriptome of 3-month-old male Heart-iKO vs. *miR-29^fl/fl^* mice (n=4 per group). b) Gene set enrichment analysis (GSEA) plot of significantly altered Molecular Signatures Database (MSigDB) datasets in each RNA-Seq experiment (upper), and selected altered datasets from WikiPathways (WP) (lower). c) MA plot of cardiomyocyte-specific transcriptome of 6-month-old male Heart-iKO (n=3) vs. *miR-29^fl/fl^* (n=2) mice. Both MA plots represent in the x-axis the log_2_ of mean normalized counts across all samples, and in the y-axis the log_2_ fold change (LFC) for each detected gene. Significant genes are highlighted in black. The top 10 genes with the highest adjusted p-values are indicated in each MA plot.

**Table 4.**
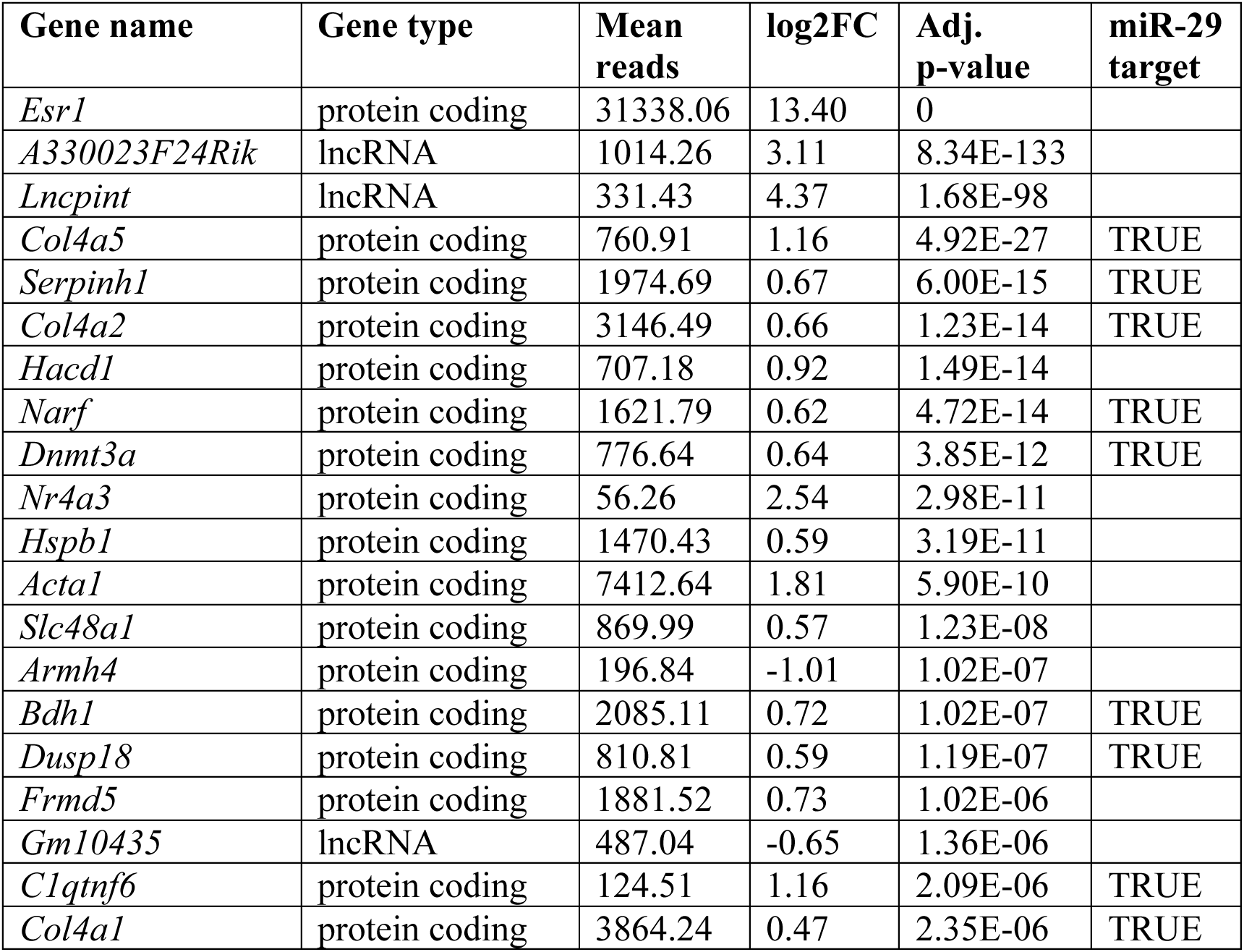
Heart-iKO bulk tissue transcriptomics. . Top differentially expressed genes in bulk heart tissue from Heart-iKO mice compared with *miR-29^fl/fl^* mice.

To further explore the biological consequences of transcriptome deregulation, we performed a GSEA on the bulk transcriptomic data, using the Hallmarks collection from the Mouse Molecular Signatures database (**Fig. 5b**). Among the most deregulated genesets, we found *oxidative phosphorylation* to be largely repressed in Heart-iKO, in agreement with our previous physiological results. We also found a repression of metabolic genesets like *fatty acid* and *xenobiotic metabolism* genes (**Fig. 5b) (Suppl. Tab 1)**. These results were in agreement with our previous results in whole-body *miR-29a/b-1^-/-^* mice, where we observed a deregulation in glucose and fatty acid metabolism pathways (12).

Given our previous observations of the specificity of miR-29 deficiency and the relevance of the non-cardiomyocyte RNA fraction in bulk heart RNA, we performed an RNA-Seq study in cardiomyocytes isolated from 6-month-old Heart-iKO and *miR-29^fl/fl^* mice to further dissociate transcriptomic changes specific to Heart-iKO cardiomyocytes. We found 179 significantly upregulated and 73 significantly downregulated genes in the cardiomyocyte Heart-iKO transcriptome, and top deregulated genes displayed a generally higher log_2_ Fold Change than those of bulk heart transcriptomics (**Fig. 5c) (Table 5**). These data supported our previous observations showing that heart transcriptomic alterations in Heart-iKO mice were essentially located in cardiomyocytes. Among the upregulated genes we found many miR-29 targets, such as those previously mentioned in the bulk heart RNA-Seq, and many others **(Suppl. Table 2)**.

**Table 5.**
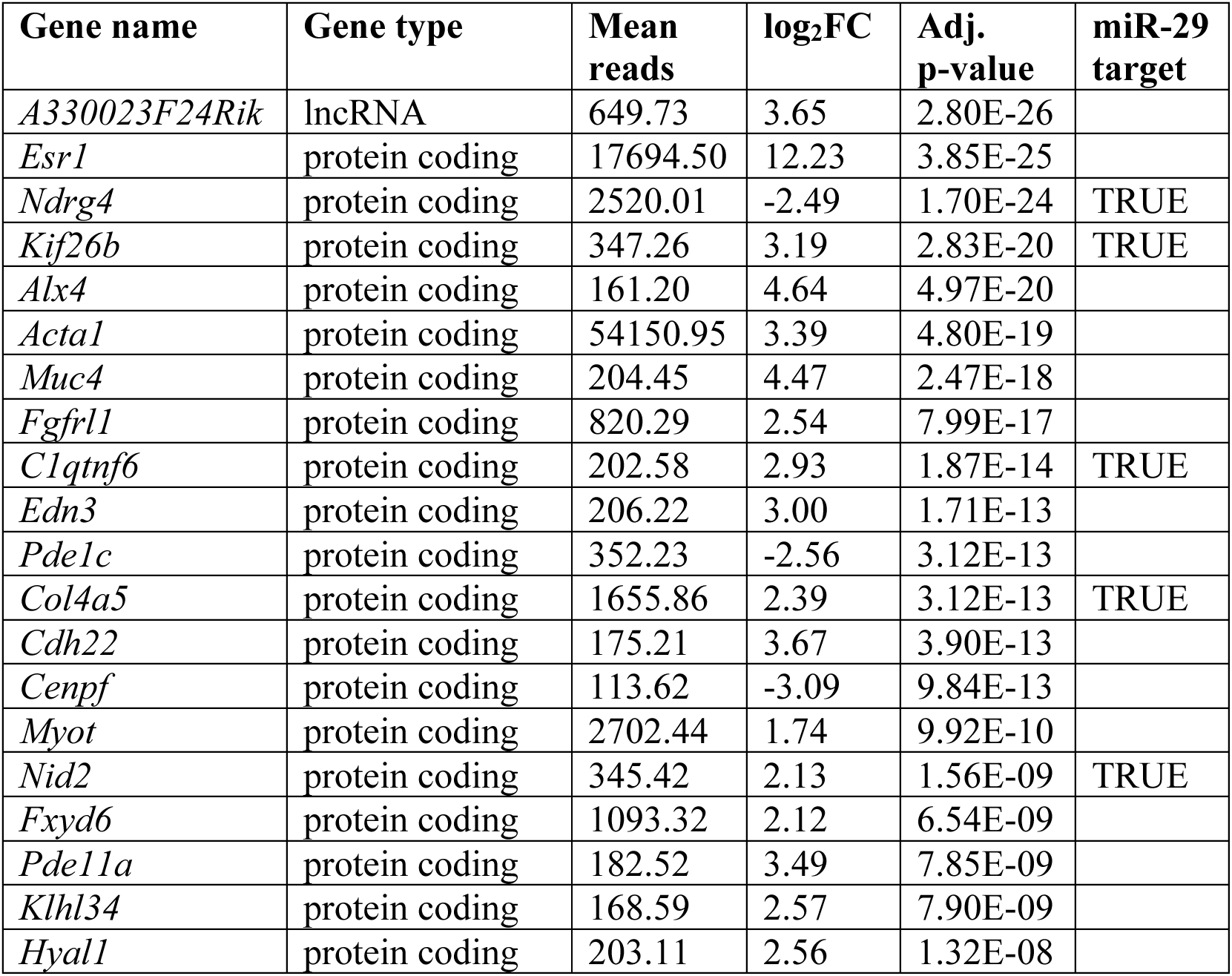
Heart-iKO cardiomyocyte transcriptomics. Top differentially expressed genes in cardiomyocytes from Heart-iKO mice compared with *miR-29^fl/fl^* mice.

Further characterization by performing a GSEA analysis found some specific alterations in Heart-iKO cardiomyocytes that were not detected in bulk heart RNA-Seq, such as the upregulation of genes related to *epithelial-mesenchymal transition* or the repression of *MYC targets* (**Fig. 5b**). On the other hand, we found a more significant repression of *oxidative phosphorylation*, *electron transport chain* genes, and *ribosomal proteins* in the Heart-iKO cardiomyocyte-specific RNA-Seq data than in the bulk heart RNA-Seq data (**Fig. 5b) (Suppl. Table 1)**.

### Cardiomyocytes derived from human iPSCs deficient in *miR-29a/b-1* display altered mitochondrial function

To elucidate the relevance of miR-29 in human cardiomyocytes, we obtained the SV10 human iPSC line from the Spanish National Bank of Stem Cell Lines and generated a *miR-29a/b-1*-deficient iPSC line by CRISPR editing. After establishing WT and *miR-29a/b-1*-deficient cell lines, we differentiated them to cardiomyocytes, obtaining CM-iPSCs. qPCR analysis of miR-29 expression displayed a full reduction of *miR-29a* levels in *miR-29a/b-1*-deficient CM-iPSCs after differentiation and 7-day maturation, as well as an almost total depletion of *miR-29b* (**Fig. 6a**). Moreover, *miR-29c* levels were reduced to around 50% of those of WT iPSCs (**Fig. 6a**). This apparent silencing of the *miR-29b-2/c* cluster could be explained by a regulatory mechanism connecting the expression of both clusters, or by the presence of basal non-specific cross-amplification of *miR-29a* in *miR-29c* qPCR as, although the TaqMan probes are copy-specific, their similarity is very high and thus cannot be discarded.

**Figure 6.**
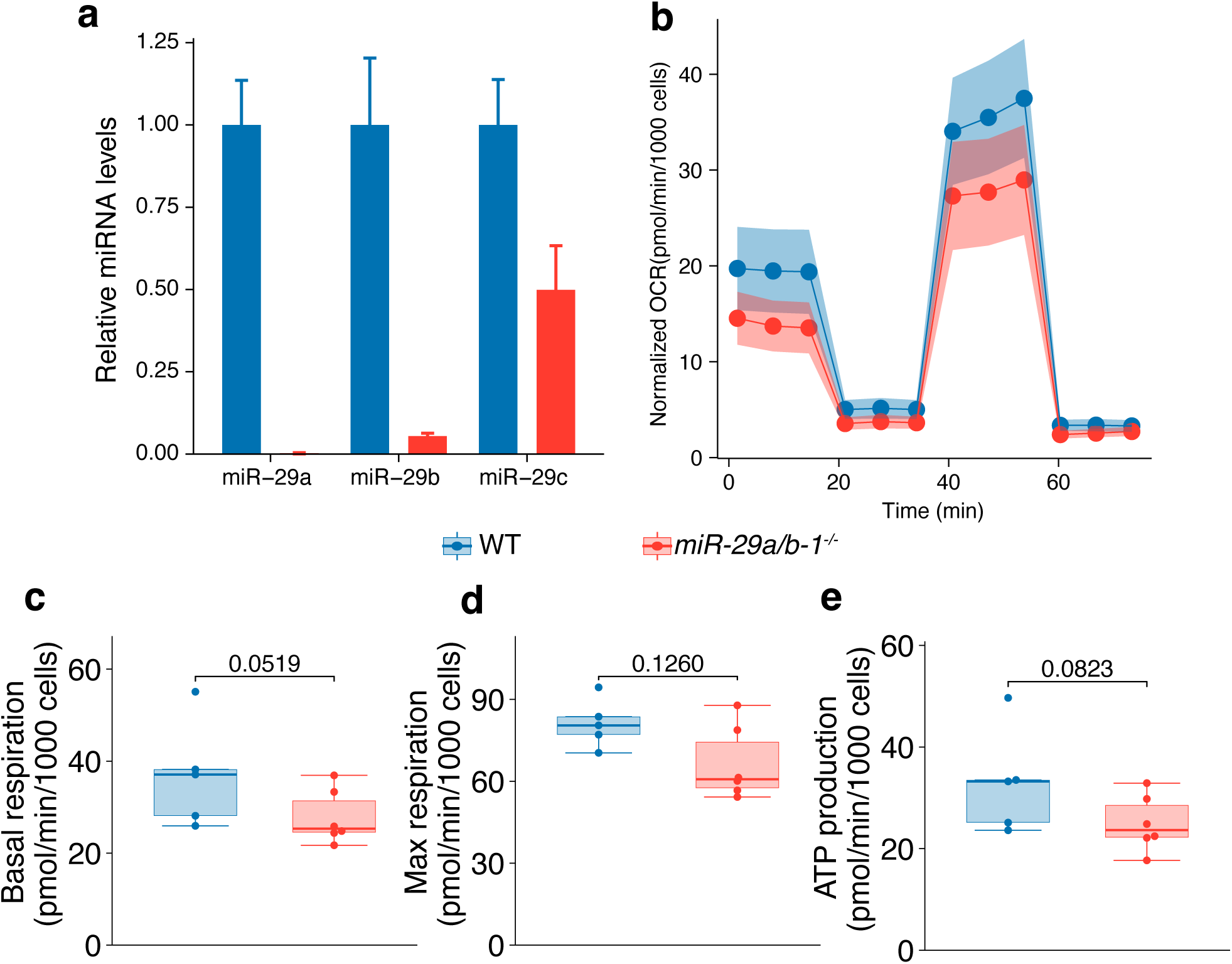
Study of miR-29 function in CM-iPSCs. a) Expression of miR-29 family members in WT and *miR-29a/b-1^-/-^* CM-iPSCs. b) Normalized oxygen consumption ratio (OCR) curve, c) basal respiration, d) maximal respiration, and e) ATP production of WT and *miR-29a/b-1^-/-^*CM-iPSCs during the Seahorse Mito Stress Test (n=5 WT and 6 *miR-29a/b-1^-/-^* replicates). Bar and line plots represent mean ± SEM. Statistical significance was determined using a non-parametric Wilcoxon signed-rank test.

After differentiation, CM-iPSCs formed a mesh with autonomous beating. There were no obvious differences in morphology between WT and *miR-29a/b-1* CM-iPSCs. To decipher whether the loss of *miR-29a/b-1* cluster also influenced CM-iPSC mitochondrial activity, we characterized mitochondrial function performing a Seahorse Mito Stress test assay (**Fig. 6b**). Consistently, *miR-29a/b-1*-deficient CM-iPSCs displayed a pattern of reduced basal and maximal respiration, as well as reduced ATP production, consistent with the defects observed in Heart-iKO mice (**Fig. 6c-e**).

## Discussion

In this study, we have established an inducible, cardiomyocyte-specific miR-29 deficient mouse model to delineate the mechanistic role of miR-29 in the adult heart. This approach allowed us to explore the cell-autonomous effects of miR-29 in cardiomyocyte function without the influence of paracrine signaling from other heart cell types, ECM remodeling or systemic influences, such as vascular alterations or complex endocrine signaling affecting heart function. By using a tamoxifen-inducible Cre recombinase, we also aimed to bypass the potentially severe developmental defects that we had previously observed in double-knockout miR-29 mice, which resulted in premature death before one month of age (12).

Our data demonstrate that miR-29 is essential for adult cardiomyocyte homeostasis, as its cardiomyocyte-specific deletion in adults caused DCM and premature death. Heart-iKO mice presented an altered heart structure and function, with an increased heart size and weight, and histology and echocardiography data also revealed an increased lumen size. Systolic function was also significantly impaired, as shown by a decrease in both EF and FS. Initial studies of miR-29 in heart function focused on its role as key regulator of cardiac fibrosis in the context of acute myocardial infarction (11). However, subsequent work by *Sassi et al.* and *Zhang et al.* demonstrated that miR-29 also plays a role in heart homeostasis independent of its function in fibroblasts (28, 29). Both studies reported a protective effect resulting from miR-29 reduction, as mice with a partial knockout of miR-29 levels were protected from transverse aortic constriction (TAC)-induced cardiac pathology. Moreover, *Zhang et al.* also found that adeno-associated virus (AAV)-mediated miR-29 overexpression in the heart provoked diastolic dysfunction (29). This apparent contradiction with our results can be explained by fundamental differences in both experimental approaches and age. Both *Sassi et al.* and *Zhang et al.* studied the response to an acutely induced pathological stress (TAC) in young animals (8-week-old), whereas our model develops a heart pathology that arises endogenously and progresses chronically over the lifespan. *Zhang et al.* study also explored the effect of miR-29 overexpression in cardiac function without additional external stressors, but their experiments are conducted at a much younger age—from 3 to 6 weeks of age. On the contrary, our Heart-iKO model has normal miR-29 levels until tamoxifen induction at 8 weeks of age. Thus, the observed differences could be explained by the distinct roles of miR-29 in development versus adult homeostasis. While reducing miR-29 levels may offer a benefit in an acute stress setting in younger animals, the long-term absence of miR-29 family is ultimately detrimental to cardiomyocyte health and survival. Furthermore, as noted previously, miR-29 expression is strongly correlated with age; therefore, its prolonged absence in adult animals may disrupt normal cardiac aging trajectories, ultimately driving pathological remodeling.

Our results with the Heart-iKO model provide additional information relative to our previous findings in the global *miR-29a/b-1^-/-^* model (12). Unlike *miR-29a/b-1^-/-^* mice, we found no abnormalities in systolic or diastolic blood pressure in Heart-iKO mice, which suggests that vascular remodeling in artery walls, and not cardiac dysfunction, was the primary mechanism behind these pressure changes. In this sense, the regulatory role of miR-29 has also been described in other vascular cell types, such as the endothelium (30), which could also exert an effect on systemic pressure. Moreover, no changes were observed in Heart-iKO mice in diastolic parameters (E/A ratio, IVRT), which could be explained by the effects of *miR-29a/b-1* deletion in non-myocytes in the whole-body knockout model, paracrine signaling from miR-29-deficient fibroblasts or changes in tissue stiffness, which could have altered diastolic behavior. Pulmonary congestion observed in *miR-29a/b-1^-/-^*mice may also be aggravated by the local deletion of *miR-29a/b-1* in lung tissue and thus could affect diastolic function. Furthermore, as miRNA functioning is dynamic and depends on the transcriptome context, the role of miR-29 might vary with the biological stage (development versus adult homeostasis and aging). Thus, the effect of miR-29 absence during development in global *miR-29a/b-1*-deficient mice could also be responsible for the divergent cardiac pathological phenotypes.

At the mechanistic level, we found mitochondrial structural changes in Heart-iKO cardiomyocytes, which were also present in *miR-29a/b-1^-/-^* mice. While the mean area of mitochondria showed no significant changes, their morphology was altered, with a notable decrease in circularity and a larger percentage of irregularly shaped mitochondria. Moreover, these organelles were also less abundant in cardiomyocytes. At the molecular level, we identified a critical defect in mitochondrial respiration in Heart-iKO mice, specifically affecting complexes I and II of the ETC. Recent work also connected miR-29 to OxPhos (31). The inhibition of miR-29 in pancreatic β-cells increased the expression of complexes I and II of the ETC and boosted mitochondrial OxPhos activity. In our Heart-iKO model, however, we found the opposite effect, as the loss of miR-29 disrupted OxPhos activity in heart mitochondria. As with other physiological processes, these findings underscore the highly pleiotropic and context-dependent role of miR-29 in regulating mitochondrial energy metabolism.

An RNA-Seq analysis further supported the mitochondrial alterations previously observed in TEM and respirometry, revealing that genesets associated with the ETC and OxPhos were strongly repressed. Furthermore, the *fatty acid metabolism* geneset was also downregulated, a finding likely related to the observed mitochondrial dysfunction and a switch in cardiomyocyte metabolism. During postnatal development, cardiomyocytes undergo a metabolic switch, partially controlled by p38γΔ kinases, shifting from glycolysis to fatty-acid-driven metabolism (32). However, certain pathological conditions can trigger a reversion to fetal-like glycolysis, a phenomenon we also observed in the *miR-29a/b-1^-/-^* whole-body KO mice (12). This pathological metabolic switch has also been linked to mitochondrial fragmentation and heart failure (33). Therefore, the interplay between metabolic dysregulation, mitochondrial dynamics, and cardiomyocyte failure appears to be a critical mechanism driving the phenotype in our model. Further work is now required to fully characterize the molecular pathways by which miR-29 orchestrates these interconnected processes.

In our search for these molecular mechanisms, we explored individual miR-29 target genes that were upregulated in our model. Among those, we found other previously described targets, such as the DNA methyltransferase *Dnmt3a* (34), the interactor with prenylated lamin A *Narf* (35), the members of the extracellular matrix *Col4a5* and *Col4a2*, and the key enzyme in the ketone oxidation pathway *Bdh1*, which has previously been described in failing heart (36). Unlike our global knockout model, where *Pgc1α* served as a clear primary driver of mitochondrial alterations, the Heart-iKO model did not present elevated levels of this gene, which suggests that *Pgc1α* overexpression in *miR-29a/b-1^-/-^*mice could be related to other heart cell types, such as fibroblasts (37), or to systemic effects. This implies that the phenotype of Heart-iKO mice has a more complex etiology that may not be driven by the alteration of a single target gene, but rather results from the cumulative disruption of a broader regulatory network. Although we identified multiple dysregulated pathways in Heart-iKO cardiomyocytes, the precise molecular mechanism linking miR-29 deficiency to mitochondrial failure requires further elucidation. Moreover, the translational implications that arise from the results of this study must be further studied using human-related models and data.

In summary, the results presented herein demonstrate the essential role of the miR-29 family in cardiomyocyte function. Specific deletion of these miRNAs in adult cardiomyocytes alters mitochondrial ultrastructure and impairs OxPhos, ultimately leading to heart failure with reduced ejection fraction, a dilated cardiomyopathy phenotype, and premature death in mice. Thus, maintaining physiological levels of miR-29 is crucial for cardiac function, with overexpression or depletion being detrimental. Other miRNAs, such as miR-208, miR-133, and miR-212/132, have been previously described as critical mediators of cardiac pathology (38–40). However, the present work offers an insight into the molecular pathways that may drive age-associated cardiac pathologies, given the remarkable and multifaceted involvement of miR-29 in the aging process itself. It is well-established that the miR-29 family is upregulated with age in both mice and non-human primates (7, 8), and its overexpression is currently studied as a therapeutic target in multiple fibrotic diseases, including heart failure (41). Our results thus serve as a cautionary note for therapeutic development, as a systemic or cardiac-specific inhibition of miR-29 could inadvertently induce mitochondrial dysfunction and heart failure.

## Acknowledgements

We thank Dr. Juan Miguel Redondo for kindly sharing *Myh6-Cre* mice. We thank A. Moyano, R. Feijóo, A. López, L. Ronderos, and D. Álvarez for technical support to this project, and Y. Español for her advice and management support. We also thank Gema Rey and Pedro M. Quirós for helpful comments and advice. This work was supported by Spanish Agencia Estatal de Investigación (AEI) (PID2020-118394RB-I00, PID2023-148089OB-I00; 10.13039/501100011033), and Consejería de Ciencia, Innovación y Universidad del Gobierno del Principado de Asturias (IDE/2024/000784). D.R.-V. was supported by a FPU fellowship from Ministerio de Ciencia y Universidades (MU-20-FPU19/00477). C.F. is supported by a Ramón y Cajal fellowship (RYC2024-050343-I) by AEI. L.M-N. is supported by the FPI program (PRE2021-100065) funded by AEI. A.G.T. is supported by a FPU fellowship from Ministerio de Ciencia, Innovación y Universidades (MCIU-24-FPU23-02504). B.C. was supported by a fellowship from “La Caixa Foundation” (ID 100010434; fellowship code LCF/BQ/DR21/11880010). R.R.-B. was supported by a FPU fellowship from Ministerio de Ciencia y Universidades (FPU17/03847). G.S. is supported by AEI (PID2022-138525OB-I00) and FEDER/EU funds, “La Caixa Foundation” (HR24-00581), ERC Synergy AdipoHealth; Comunidad de Madrid (Syg-2024/Salgl-1012, AdipohearTCM), and “Cris Contra el Cancer” (PR_EX_2024-22). X.M.C. is supported by an AEI grant (PID2024-159805NA-I00) and AEI Ramón y Cajal fellowship (RYC2023-042853-I). A.P.U. is supported by a Ramón y Cajal fellowship (RYC2021-031776-I) by AEI.

## Author contributions

D.R.-V., C.L.-O., X.M.C., and A.P.U. designed research; G.S. designed cardiovascular research protocols; Y.-W.H. designed and generated *miR-29 floxed* mice; D.R.-V. conducted research; C.F., B.C., and R.R.-B. contributed to cardiovascular characterization; L.M.-N., A.G.T., and F.R. contributed to mouse experimental work; D.R.-V., J.M.P.F., C.L.-O., X.M.C., and A.P.U. wrote the manuscript, with input from all authors. All the authors approved the submitted version of the manuscript.

---

**Supplementary Figure 1.**
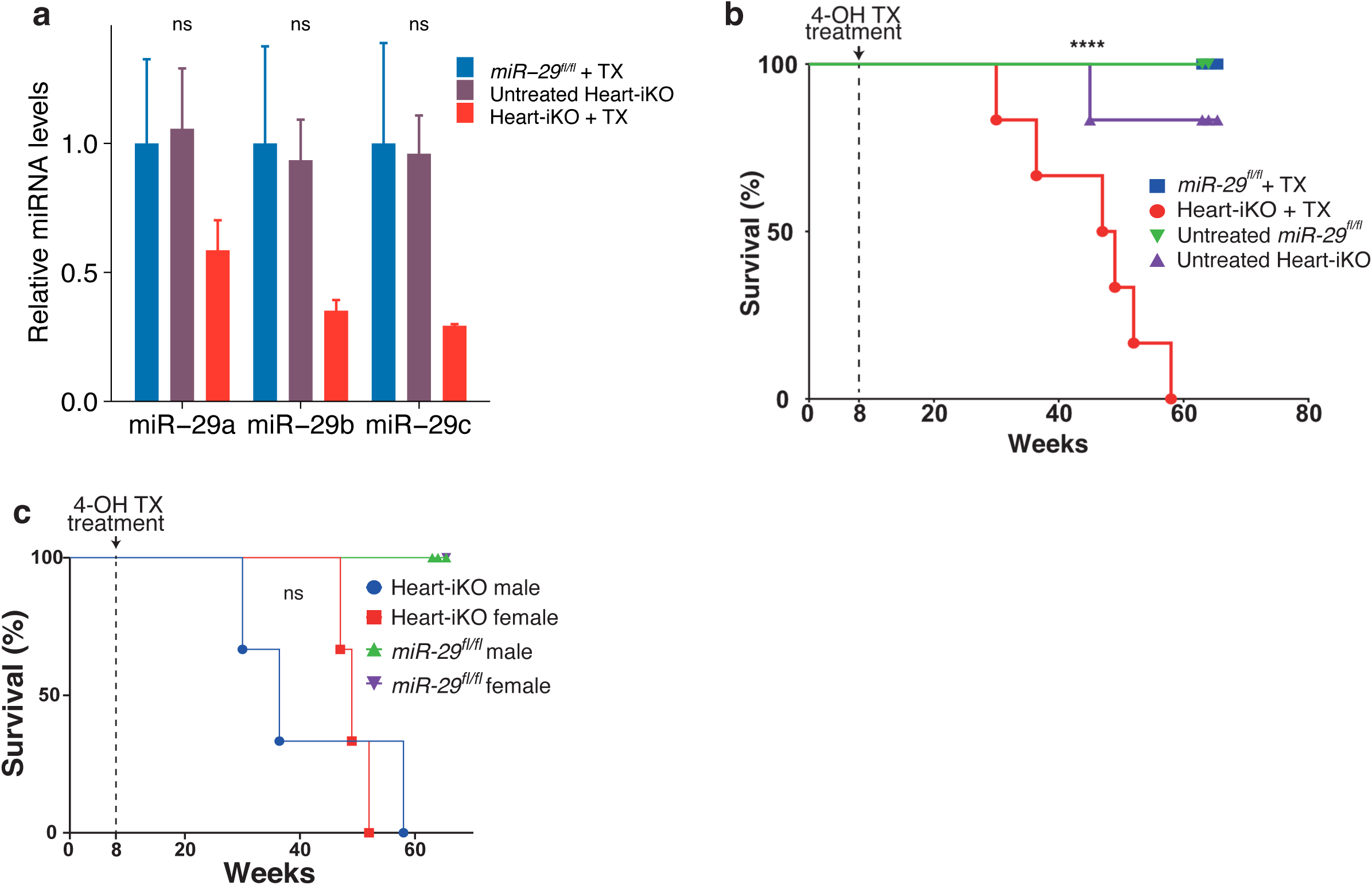
Characterization of Heart-iKO mice. a) Expression levels of each miR-29 family member in heart tissue from *miR-29^fl/fl^* mice treated with tamoxifen, untreated Heart-iKO and tamoxifen-treated Heart-iKO mice (n=2 mice per group, 1 month after treatment). b) Survival curve of untreated *miR-29^fl/fl^*mice (n=6), *miR-29a/b-1^-/-^*mice treated with tamoxifen (TX) (n=7), untreated Heart-iKO (n=4), and tamoxifen-treated Heart-iKO mice (n=6). c) Survival curve divided by sex of tamoxifen-treated *miR-29^fl/fl^* (n=6 male and 1 female) and Heart-iKO (n=3 male and 3 female) mice. Bar and weight line plots represent mean ± SEM. Statistical significance between male and female Heart-iKO was determined by a log-rank test in the survival analysis. In the relative expression data, statistical significance was calculated using a two-way ANOVA and a post-hoc Tukey analysis. *: p < 0.05, **: p < 0.01, ***: p < 0.001, ****: p < 0.0001, ns: p > 0.05.

**Supplementary Figure 2.**
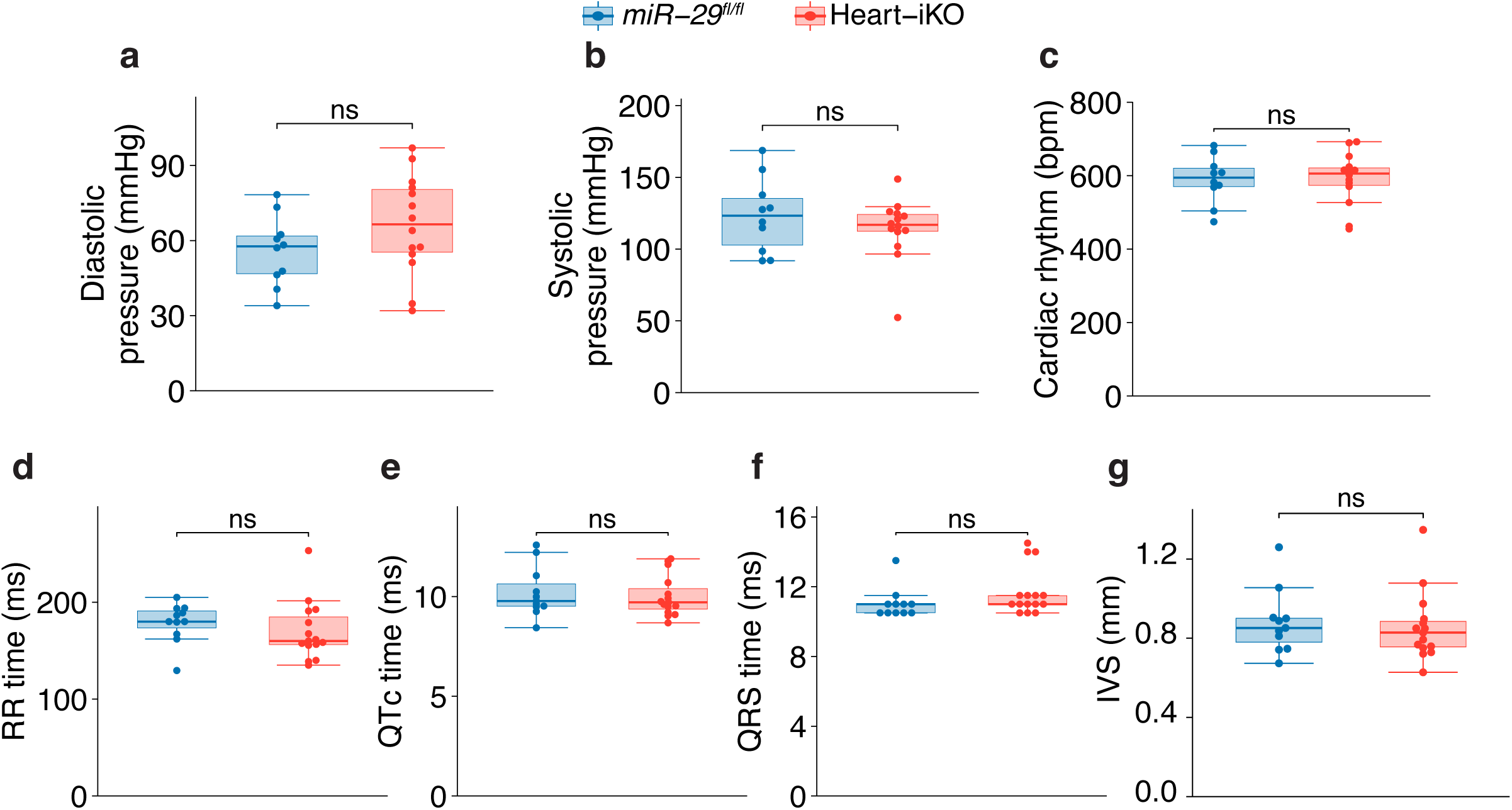
Blood pressure analysis, electrocardiography, and additional echocardiographic analysis of Heart-iKO mice. a) Diastolic pressure, b) systolic pressure, and c) cardiac rhythm of 5-month-old male *miR-29^fl/fl^* (n=11) and Heart-iKO (n=15) mice. d) RR time, e) QTc time, and f) QRS time electrocardiogram parameters of 5-month-old male *miR-29^fl/fl^* (n=11) and Heart-iKO (n=15) mice. g) Diastolic intraventricular septum (IVS) length of 5-month-old male *miR-29^fl/fl^* (n=11) and Heart-iKO (n=15) mice. Statistical significance was determined using a non-parametric Wilcoxon signed-rank test. ns: p > 0.05.

**Supplementary Figure 3.**
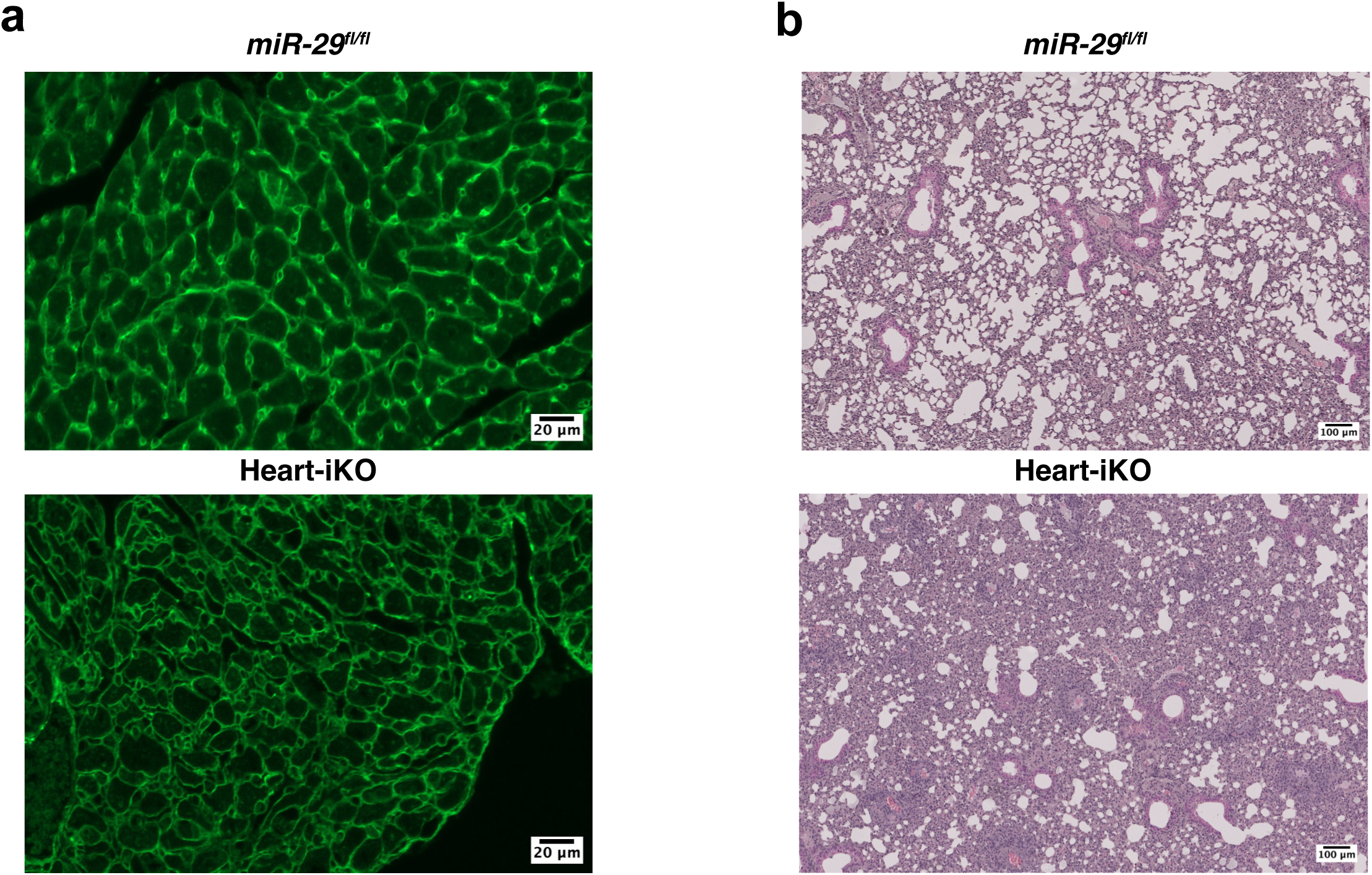
Macroscopic heart images and additional histological characterization of Heart-iKO mice. a) WGA-stained tissue section, and b) lung H/E-stained tissue section of a 6-month-old control *miR-29^fl/fl^* compared to an advanced DCM in a Heart-iKO mice of the same age.

**Supplementary Figure 4.**
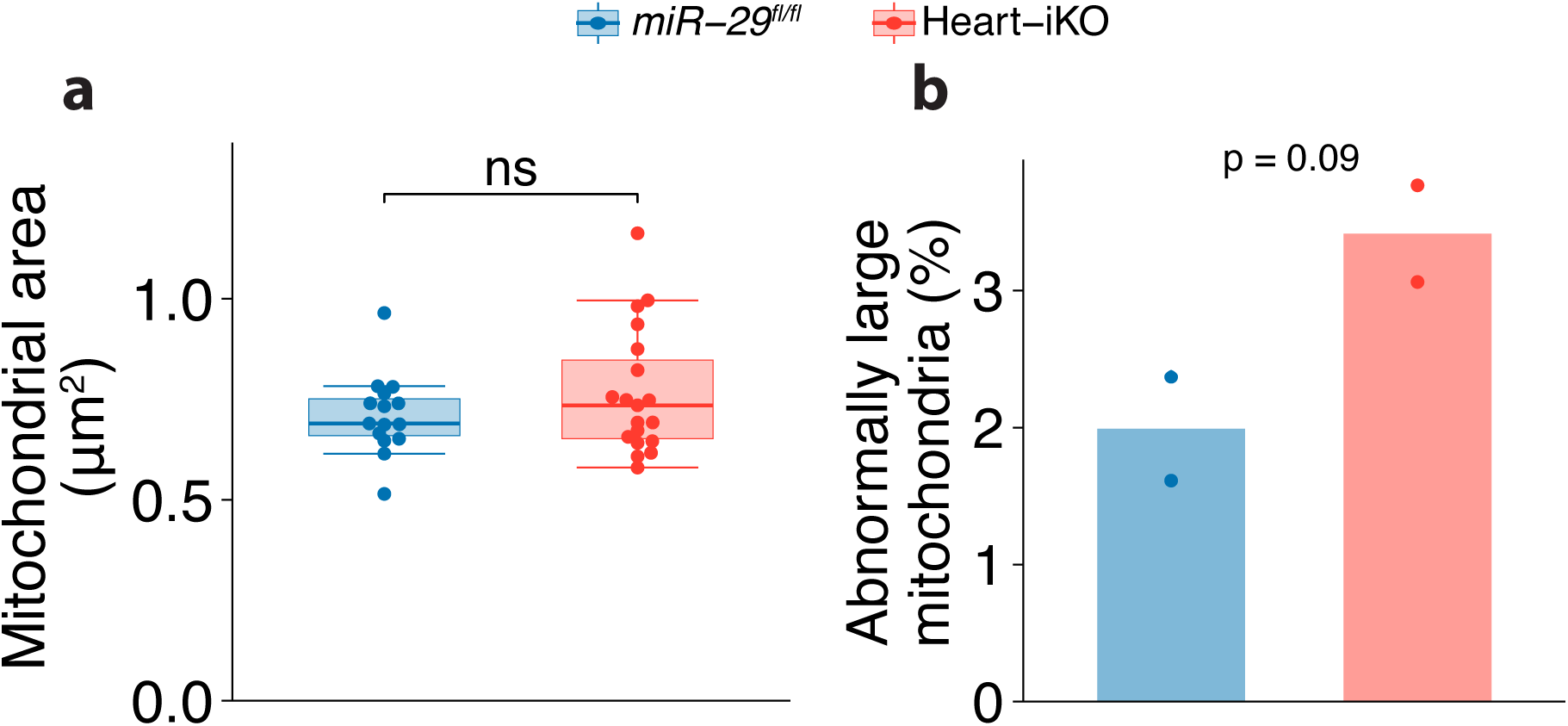
Additional mitochondrial ultrastructural parameters of Heart-iKO mice. a) Mitochondrial area, and b) percentage of mitochondria with an abnormally large area (> 2 µm^2^) in cardiomyocytes from 5-month-old Heart-iKO and *miR-29^fl/fl^* mice. Images were obtained from 5 tissue slices from 2 male mice per genotype. Each data point represents one image (n=15 *miR-29^fl/fl^*, 19 Heart-iKO). Bar plots represent mean values. Statistical significance was determined using a non-parametric Wilcoxon signed-rank test for area measurements. For the percentage of abnormally large mitochondria, data was structured as a contingency table and analyzed using a chi-squared test. ns: p > 0.05.

**Supplementary Table 1.** Leading-edge genes from highlighted genesets in Heart-iKO. Top 30 leading-edge genes from highlighted genesets in the GSEA of Heart-iKO transcriptomics.

| Data | Pathway | Leading-edge genes |
| --- | --- | --- |
| CM spec. | OxPhos (MSigDB) | *Etfb, Slc25a5, Ndufs6, Cox7a2, Cox17, Atp5b, Atp5e, Cox7b, Atp5h, Surf1, Cox5b, Immt, Atp5k, Atp5g1, Ndufs7, Cox7c, Ndufs2, Ndufb7, Cox5a, Ndufa5, Cox6a1, Mdh2, Atp6v1d, Suclg1, Cox8a, Ndufs8, Slc25a11, Ndufa3, Uqcrc1, Ndufv1* |
| CM spec. | ETC (WP) | *Cox7a1, mt-Nd3, Slc25a5, Ndufs6, Cox7a2, Ndufa12, Cox17, Ndufv3, Cox7b, Surf1, Cox5b, Ndufs7, Ndufs2, Ndufb7, Cox5a, Ndufa5, Cox6a1, Cox8a, Ndufs8, Ndufb10, Uqcrc1, Ndufv1, Cox4i1, Ndufs5, Ndufv2, Cox6c, Uqcr11, Ndufa8, Ndufb9, Ndufab1* |
| CM spec. | Cytoplasmic ribosomal proteins (WP) | *Rpl38, Rps8, Rpl28, Rps16, Rps17, Rpl13a, Rps21, Fau, Rpl37a, Rps2, Rps6, Rps15, Rps19, Rps23, Rps20, Rpl8, Rpl21, Rpsa, Rpl15, Rplp1, Rpl7a, Rpl18a, Rpl35, Rpl17, Rpl19, Rpl13, Rpl41, Rps3, Rpl10, Rpl10a* |
| CM spec. | OxPhos (WP) | *Ndufa4l2, mt-Nd3, Ndufs6, Ndufv3, Ndufs7, Ndufs2, Ndufb7, Ndufa5, Ndufs8, Ndufb10, Ndufv1, Ndufb8, Ndufs5, Ndufv2, Ndufa8, Ndufa11, Ndufb9, Ndufa7, Ndufb2, Ndufa2, Ndufa9, Ndufc1, Atp5pb, Ndufs4, Ndufa6, Ndufa10, mt-Nd6, Ndufb5* |
| Bulk | ETC (WP) | *mt-Nd6, Cox6c, Ndufs2, Cox17, mt-Co2, mt-Nd1, Uqcr11, Ucp3, Ndufa12, Ndufs8, Uqcrc2, Cox7a2, Sdha, Ndufv3, mt-Co3, mt-Co1, mt-Nd5, Slc25a14, Uqcrh, Ndufb10, mt-Nd2, Ndufs6, Ndufs1, Cox6a2, Slc25a5, mt-Cytb, Ndufs7, Ndufc2, Ndufv1, Ndufa9* |
| Bulk | OxPhos (MSigDB) | *Nqo2, Retsat, Maob, Pdp1, Cyc1, Cox6c, Ndufs2, Cox17, Idh3g, Alas1, Hadhb, Idh1, Etfb, Uqcr11, Ech1, Acaa2, Suclg1, Mdh1, Atp5h, Pdha1, Ndufs8, Acadvl, Decr1, Uqcrc2, Cox7a2, Cpt1a, Sdha, Etfdh, Dld, Cox7c* |
| Bulk | Cytoplasmic ribosomal proteins (WP) | *Gm15501, Rpl26, Fau, Rps5, Rpl7a, Rpl30, Rpl23a, Rplp1, Rps2, Rpl13, Rpl18a, Rps12, Rpl15, Rps18, Rplp0, Rps14, Rps21, Rpl12, Rpl34, Rps17, Rps10, Rpl7, Rps6, Rps24, Rpl13a, Rpl17, Rpl31, Rpl11, Rpl41, Rpl6* |
| Bulk | OxPhos (WP) | *mt-Nd6, Ndufs2, mt-Nd1, Ndufs8, Ndufv3, Ndufb8, mt-Nd5, Ndufb10, mt-Nd2, Ndufs6, Ndufs1, Ndufs7, Ndufc2, Ndufv1, Ndufa9, Ndufa6, mt-Nd4, Ndufb7, mt-Atp6, Ndufa7, Ndufa5, Ndufb2, Atp5pb, Ndufb9, mt-Nd3, Ndufv2, Ndufc1, Ndufa11* |

**Supplementary Table 2.**
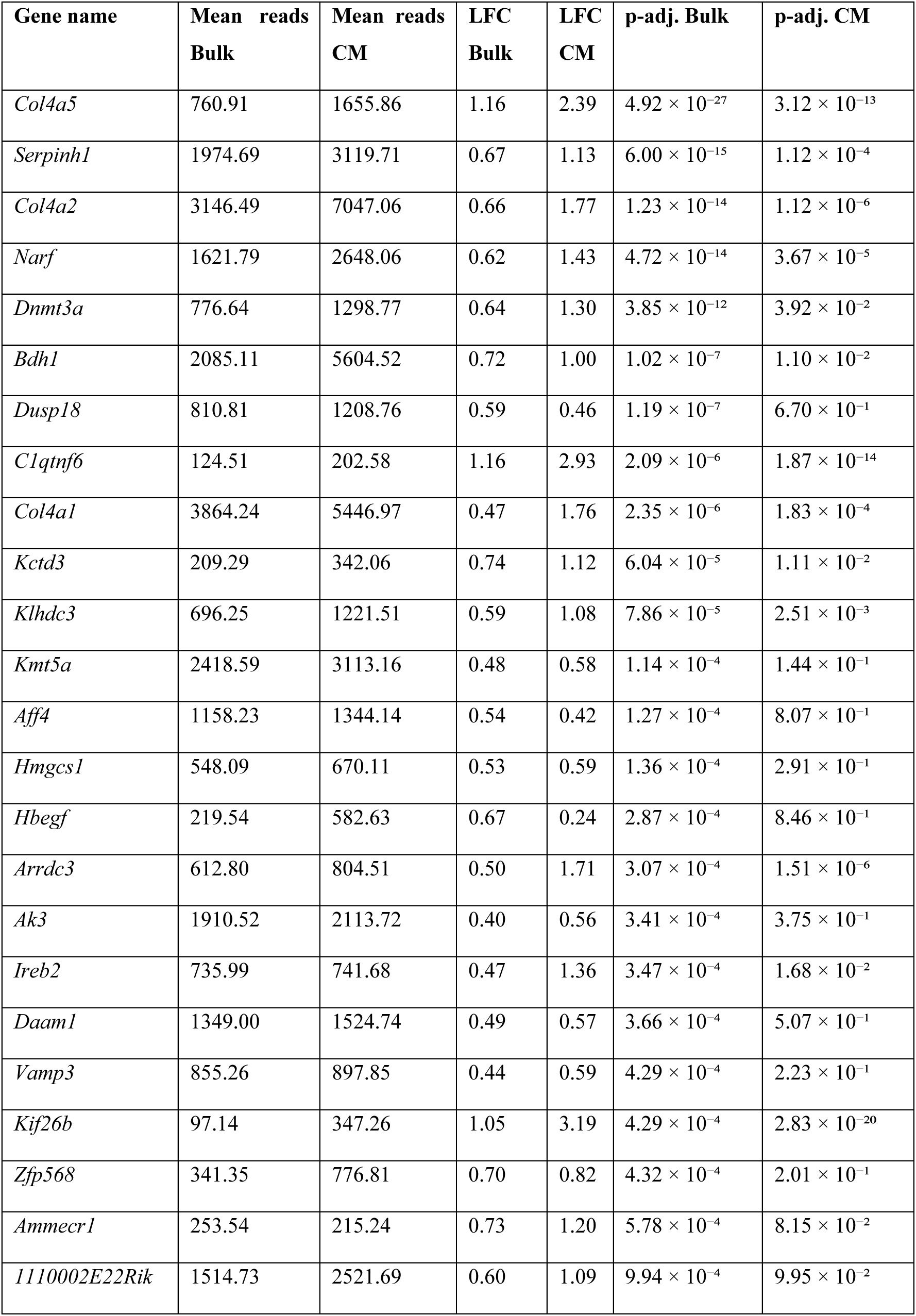
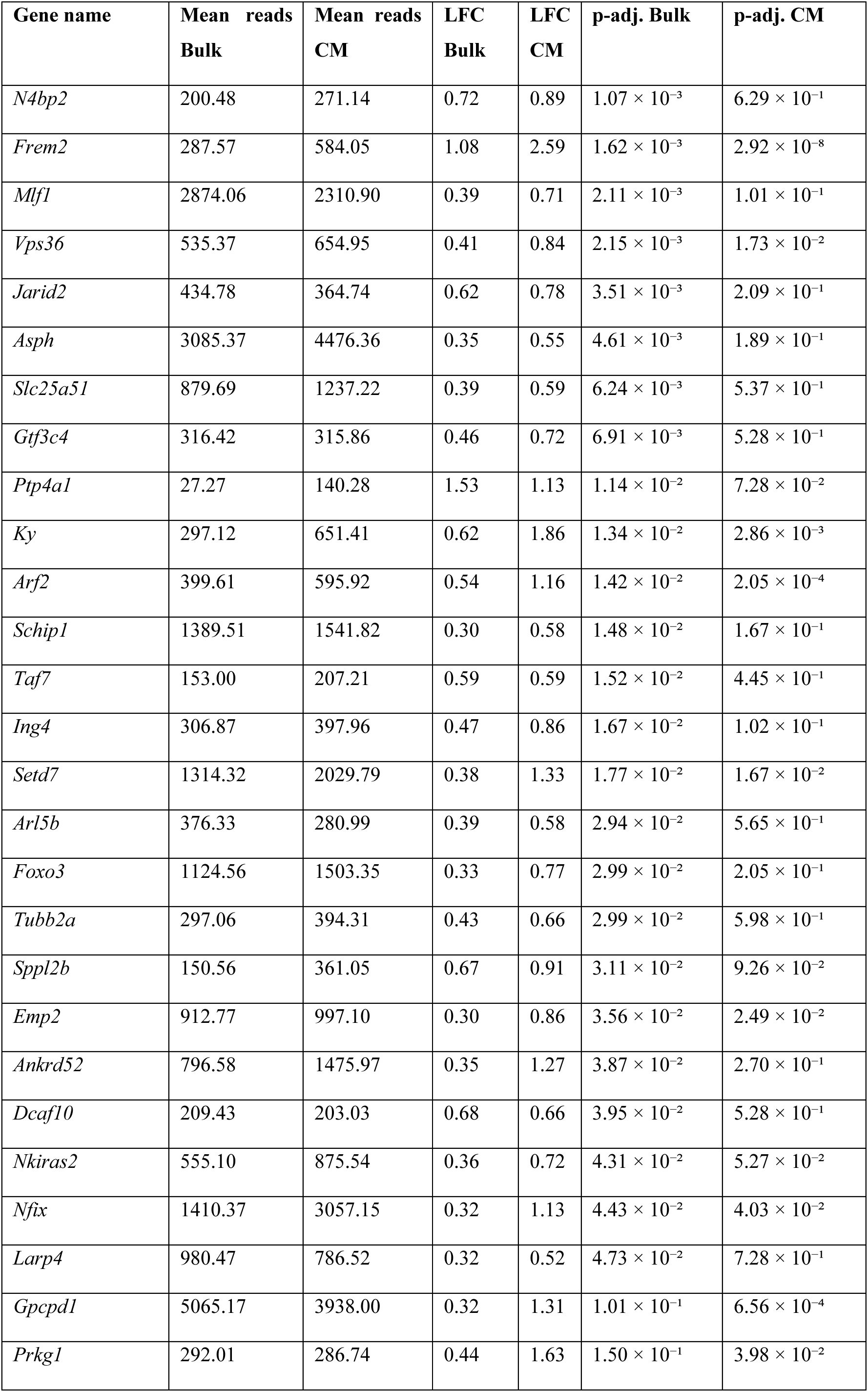
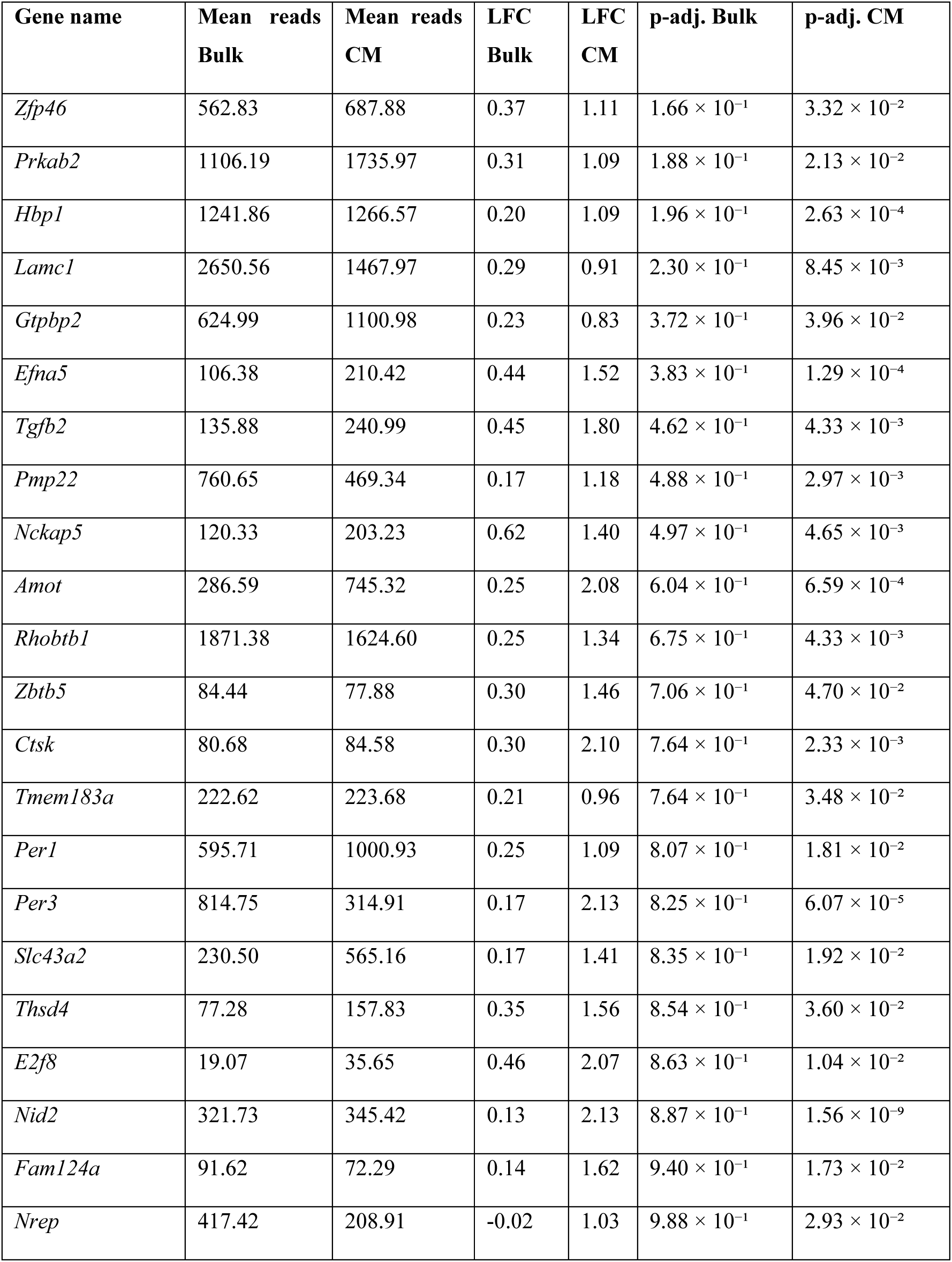
miR-29 targets upregulated in Heart-iKO transcriptomic analyses. Genes encoding miR-29 target transcripts that are upregulated in either bulk or cardiomyocyte-specific Heart-iKO transcriptomics with a LFC > 0.5.

## References

1. B. K. Kennedy, et al., Geroscience: Linking Aging to Chronic Disease. Cell 159, 709–713 (2014).

2. M. Naghavi, et al., Global burden of 288 causes of death and life expectancy decomposition in 204 countries and territories and 811 subnational locations, 1990–2021: a systematic analysis for the Global Burden of Disease Study 2021. The Lancet 403, 2100–2132 (2024).

3. C. López-Otín, M. A. Blasco, L. Partridge, M. Serrano, G. Kroemer, Hallmarks of aging: An expanding universe. Cell 186, 243–278 (2023).

4. G. Kroemer, et al., From geroscience to precision geromedicine: Understanding and managing aging. Cell 188, 2043–2062 (2025).

5. M. Abdellatif, P. P. Rainer, S. Sedej, G. Kroemer, Hallmarks of cardiovascular ageing. Nat. Rev. Cardiol. 1–24 (2023). 10.1038/s41569-023-00881-3.

6. A. P. Ugalde, et al., Aging and chronic DNA damage response activate a regulatory pathway involving miR-29 and p53: Regulatory circuitry involving miR-29 and p53. EMBO J. 30, 2219–2232 (2011).

7. V. Wagner, et al., Characterizing expression changes in noncoding RNAs during aging and heterochronic parabiosis across mouse tissues. Nat. Biotechnol. 42, 109–118 (2024).

8. V. Swahari, et al., miR-29 is an important driver of aging-related phenotypes. Commun. Biol. 7, 1055 (2024).

9. J. J. Kwon, T. D. Factora, S. Dey, J. Kota, A systematic review of miR-29 in cancer. Mol. Ther. Oncolytics 12, 173–194 (2019).

10. J. Massart, et al., Altered miR-29 expression in type 2 diabetes influences glucose and lipid metabolism in skeletal muscle. Diabetes 66, 1807–1818 (2017).

11. E. Van Rooij, et al., Dysregulation of microRNAs after myocardial infarction reveals a role of miR-29 in cardiac fibrosis. Proc. Natl. Acad. Sci. 105, 13027–13032 (2008).

12. X. M. Caravia, et al., The microRNA-29/PGC1α regulatory axis is critical for metabolic control of cardiac function. PLOS Biol. 16, e2006247 (2018).

13. D. S. Sohal, et al., Temporally regulated and tissue-specific gene manipulations in the adult and embryonic heart using a tamoxifen-inducible cre protein. Circ. Res. 89, 20–25 (2001).

14. D. Li, J. Wu, Y. Bai, X. Zhao, L. Liu, Isolation and culture of adult mouse cardiomyocytes for cell signaling and in vitro cardiac hypertrophy. J. Vis. Exp. 51357 (2014). 10.3791/51357.

15. X. M. Caravia, et al., Precise gene editing of pathogenic lamin a mutations corrects cardiac disease. Proc. Natl. Acad. Sci. U. S. A. 122, e2515267122 (2025).

16. D. A. M. Feyen, et al., Metabolic maturation media improve physiological function of human iPSC-derived cardiomyocytes. Cell Rep. 32, 107925 (2020).

17. Y. Kubota, et al., Evaluation of blood pressure measured by tail-cuff methods (without heating) in spontaneously hypertensive rats. Biol. Pharm. Bull. 29, 1756–1758 (2006).

18. R. Romero-Becerra, et al., p38γ/δ activation alters cardiac electrical activity and predisposes to ventricular arrhythmia. *Nat*. Cardiovasc. Res. 2, 1204–1220 (2023).

19. R. Conrad, K. Narayan, Instance segmentation of mitochondria in electron microscopy images with a generalist deep learning model trained on a diverse dataset. Cell Syst. 14, 58–71.e5 (2023).

20. J. B. Engler, Tidyplots empowers life scientists with easy code-based data visualization. iMeta 4, e70018 (2025).

21. M. I. Love, W. Huber, S. Anders, Moderated estimation of fold change and dispersion for RNA-seq data with DESeq2. Genome Biol. 15, 550 (2014).

22. A. S. Castanza, et al., Extending support for mouse data in the Molecular Signatures Database (MSigDB). Nat. Methods 20, 1619–1620 (2023).

23. G. Korotkevich, et al., Fast gene set enrichment analysis. [Preprint] (2021). Available at: https://www.biorxiv.org/content/10.1101/060012v3 [Accessed 23 May 2025].

24. A. Kuznetsova, P. B. Brockhoff, R. H. B. Christensen, lmerTest Package: Tests in Linear Mixed Effects Models. J. Stat. Softw. 82, 1–26 (2017).

25. T. M. Therneau, P. M. Grambsch, Modeling survival data: extending the cox model (Springer, 2000).

26. T. Nishiyama, et al., Precise genomic editing of pathogenic mutations in RBM20 rescues dilated cardiomyopathy. Sci. Transl. Med. 14, eade1633 (2022).

27. E. Gnaiger, B. Lassnig, A. Kuznetsov, G. Rieger, R. Margreiter, Mitochondrial oxygen affinity, respiratory flux control and excess capacity of cytochrome c oxidase. J. Exp. Biol. 201, 1129–1139 (1998).

28. Y. Sassi, et al., Cardiac myocyte miR-29 promotes pathological remodeling of the heart by activating Wnt signaling. Nat. Commun. 8, 1614 (2017).

29. X. Zhang, et al., Modulation of miR-29 influences myocardial compliance likely through coordinated regulation of calcium handling and extracellular matrix. Mol. Ther. Nucleic Acids 34, 102081 (2023).

30. M. E. Widlansky, et al., miR-29 contributes to normal endothelial function and can restore it in cardiometabolic disorders. EMBO Mol. Med. 10, EMMM201708046 (2018).

31. E. Cowan, et al., MicroRNA 29 modulates β-cell mitochondrial metabolism and insulin secretion via underlying miR-29-OXPHOS complex pathways. Acta Physiol. 240, e14180 (2024).

32. A. M. Santamans, et al., p38γ and p38δ regulate postnatal cardiac metabolism through glycogen synthase 1. PLOS Biol. 19, e3001447 (2021).

33. T. Wai, et al., Imbalanced OPA1 processing and mitochondrial fragmentation cause heart failure in mice. Science (2015). 10.1126/science.aad0116.

34. M. Fabbri, et al., MicroRNA-29 family reverts aberrant methylation in lung cancer by targeting DNA methyltransferases 3A and 3B. Proc. Natl. Acad. Sci. 104, 15805–15810 (2007).

35. R. M. Barton, H. J. Worman, Prenylated prelamin A interacts with Narf, a novel nuclear protein. J. Biol. Chem. 274, 30008–30018 (1999).

36. G. Aubert, et al., The failing heart relies on ketone bodies as a fuel. Circulation 133, 698–705 (2016).

37. H. Chen, X. Pan, L. Ding, C. Ruan, P. Gao, Cardiac fibroblast-specific knockout of PGC-1α accelerates AngII-induced cardiac remodeling. Front. Cardiovasc. Med. 8, 664626 (2021).

38. A. Carè, et al., MicroRNA-133 controls cardiac hypertrophy. Nat. Med. 13, 613–618 (2007).

39. E. Van Rooij, et al., Control of stress-dependent cardiac growth and gene expression by a microRNA. Science 316, 575–579 (2007).

40. A. Ucar, et al., The miRNA-212/132 family regulates both cardiac hypertrophy and cardiomyocyte autophagy. Nat. Commun. 3, 1078 (2012).

41. X. Wang, I. Anwar, C. P. Hodgkinson, miRNAs to the rescue: reversing heart failure by targeting miR-29. Mol. Ther. Nucleic Acids 35, 102105 (2024).

